# Archetype analysis of lung adenocarcinoma premalignancy links heterogeneity in premalignant lesions to diverging features of invasive disease

**DOI:** 10.1101/2025.10.04.680402

**Authors:** Kelley E. Anderson, Linh M. Tran, Kostyantyn Krysan, Yohana Kefella, Lingge Yu, Gregory A. Fishbein, Erika Rodriguez, Maryam Shabihkhani, Daniel P. Stefanko, Emily Green, Gang Liu, Hanqiao Liu, Sherry Zhang, Erin Kane, Mitra Mehrad, Avrum E. Spira, Steven M. Dubinett, Eric J. Burks, Sarah A. Mazzilli, Marc E. Lenburg, Jennifer E. Beane

## Abstract

Lung adenocarcinoma (**LUAD**), the most common form of lung cancer, is a heterogeneous disease with highly variable histopathologic features and clinical outcomes. LUAD premalignant lesions (**PMLs**) are localized proliferative lesions of atypical pneumocytes that proliferate along the alveolar walls. In this study, we characterized the molecular heterogeneity across PMLs, regardless of histologic designation, and identified four unique groups or “archetypes” of PMLs using bulk RNA and exome sequencing data from laser captured microdissected PMLs, normal, and tumor tissue from LUAD resection cases. The archetypes were defined based on expression of recurrent gene co-expression modules discovered using the LUAD PMLs we profiled and three publicly available datasets. One PML archetype, termed “proliferation”, was associated with increased expression of genes involved in cell proliferation, a pro-tumor immune environment, and enrichment for *EGFR* driver mutations. Another archetype, termed “normal-like”, was associated with similar features to normal tissue, an anti-tumor immune response, and lacked enrichment for driver mutations compared to other archetypes. We projected the PML archetypes into independent gene expression datasets profiling LUAD and found that tumors closest to the proliferation archetype were enriched for multiple features of aggressive LUAD, including high tumor grade and lymph node invasion, and had significantly lower disease-free survival. Proliferation archetype PMLs may represent a subset of lesions with more aggressive features that could improve patient stratification and suggest novel targets for lung cancer interception.

## INTRODUCTION

Lung cancer is the most common cause of cancer-related death, both worldwide^1^ and in the USA, with an overall five-year relative survival of 26%^2^. The most common form of lung cancer is adenocarcinoma (**LUAD**). LUAD has highly variable clinical outcomes and is heterogenous in terms of histopathologic features and molecular alterations. LUAD premalignant lesions (**PMLs**) are precursors to LUAD and include adenomatous hyperplasia (**AAH**, a localized proliferation of atypical type II pneumocytes ≤ 5 mm), adenocarcinoma in situ (**AIS**, proliferation with pure lepidic growth ≤ 30 mm without invasion or necrosis), and minimally invasive adenocarcinoma (**MIA**, predominantly lepidic growth measuring ≤ 30 mm and with ≤ 5 mm of a non-lepidic component or stromal invasion)^3^. AAH and AIS/MIA typically appear as ground glass opacities on lung computed tomography (**CT**) scans, and approximately 20% of these lesions will progress to invasive malignancy^4^. With the expanding adoption of lung cancer screening programs, the detection of PMLs via lung CT is anticipated to rise substantially^5^ and advanced robotic bronchoscopy techniques are beginning to provide an opportunity to biopsy these lesions. The clinical management of PMLs, however, remains a key challenge. Image and molecular biomarkers may be useful to accurately stratify the risk of lesion progression to invasive disease^6^. Biomarkers capable of preoperatively identifying indolent preinvasive LUAD have the potential to mitigate overtreatment of screen-detected nodules^7^, while biomarkers specific for aggressive PMLs will ensure appropriate intervention.

Prior work has identified molecular features associated with premalignant and early-stage LUAD histologic progression, including somatic mutations^8–13^, transcriptional alterations^10,14–17^, and altered immune responses^11–14^. However, recent evidence from a study of LUAD premalignancy in a cohort of East Asian never smokers suggests that gene expression changes do not follow a linear progression from AAH to AIS to MIA during carcinogensis^18^, indicating that molecular changes among PMLs may not directly correspond to histologic progression. While many of the prior studies contained limited PML samples, they highlight the potential to enhance histopathological grading of PMLs by integrating molecular features. Archetype analysis has been proposed as a framework for understanding tumor heterogeneity^19,20^, and has been used to discover new targets against small cell lung cancer’s aggressive features^21^. To better understand heterogeneity among LUAD PMLs, we defined archetypes through analysis of bulk mRNA sequencing of PMLs collected from a retrospective cohort of patients and public datasets. Archetyping identified heterogeneity in gene expression patterns and pathway activity among LUAD PMLs, independent of PML histology. Using whole exome sequencing and multiplex immunofluorescence, we linked the archetypes to driver mutations and features of genomic instability, as well as to altered density of immune cell types. Projection of the archetypes into independent LUAD gene expression datasets demonstrated their association with LUAD histopathologic features and disease-free survival. Characterizing PML archetypes may enable identification of aggressive PMLs prior to their progression to invasive LUAD, as well as identify indolent PMLs that can be monitored via screening.

## RESULTS

### Study population

Tissue blocks were obtained from a retrospective cohort of patients, collected as part of the Human Tumor Atlas Network (**HTAN**) Lung Pre-Cancer Atlas (**PCA**), that underwent surgical LUAD resection at Boston Medical Center, Vanderbilt University Medical Center, and University of California, Los Angeles Health (**Table 1**). RNA and DNA from multiple lesions (PML or tumor) and normal tissue per patient were isolated from fixed tissue sections by laser-capture microdissection. RNA was subjected to bulk RNA sequencing (n=177 total samples from n=40 total patients, with n=80 AAH and AIS/MIA samples from n=29 patients, **Figure 1A**). Samples were predominantly early-stage LUAD (80% stage I) from older patients (mean age 68.7 +/- 9.6) with a history of tobacco use (62.5 % former smokers). A representative subset (see *Methods*) of these tissue samples underwent exome sequencing (n=90 LUAD and PMLs from n=25 patients), using paired normal samples from each patient as the respective germline control.

**Figure 1:**
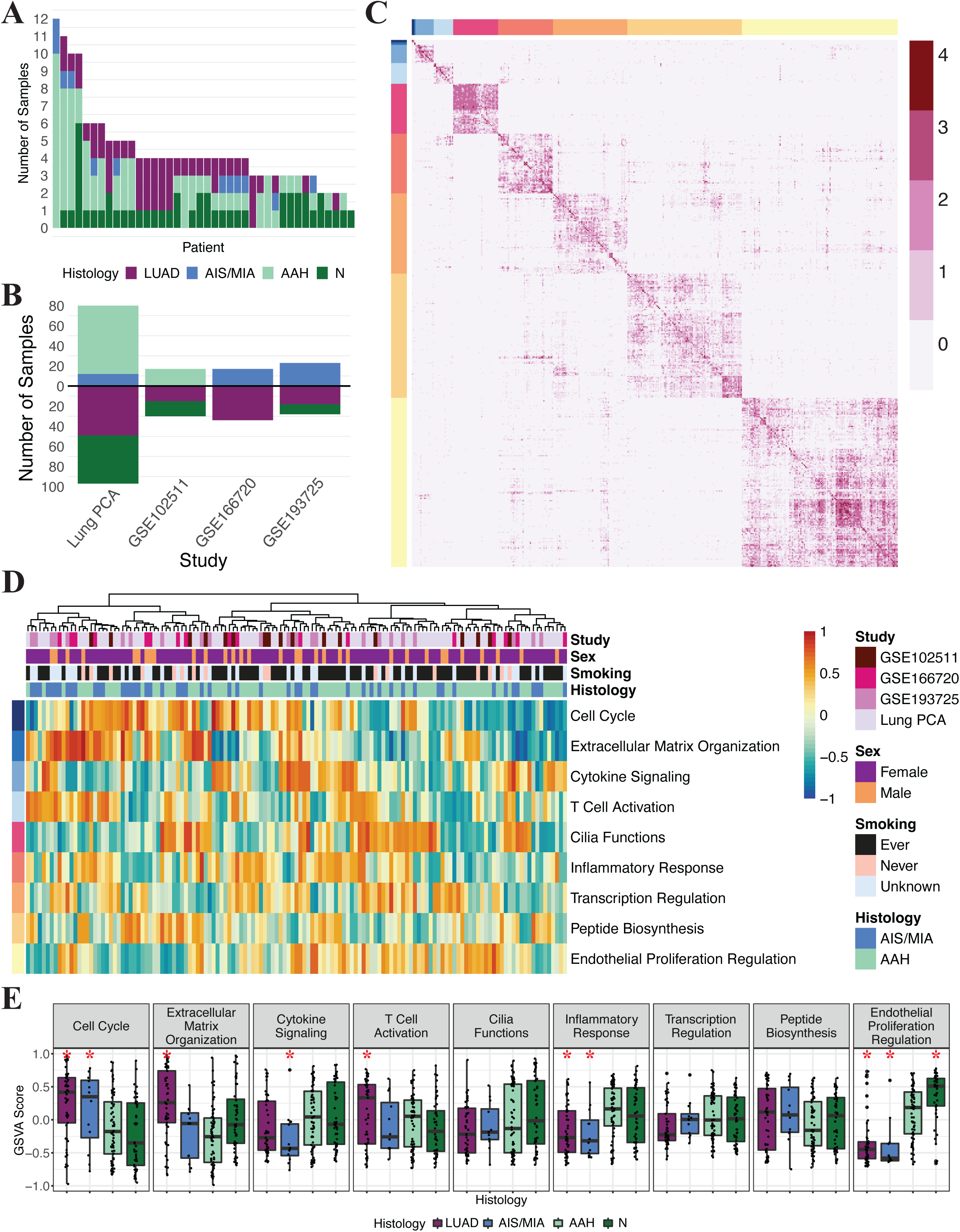
Gene modules recur across studies of LUAD premalignancy. (A) Stacked bar plot of sample histology per patient from Lung PCA cohort (n=40 patients, n=177 samples following filtering) (B) Stacked bar plot showing the histology distribution for each cohort utilized in this analysis. The cohorts included were the Lung PCA (n=80 PML, n=48 normal samples, n=49 tumor samples), GSE102511 which contained paired normal/PML/tumor samples from 17 patients (n=17 PML, n=15 normal samples, n=15 tumor samples), GSE166720 which contained independent samples from 50 patients with either early stage LUAD or MIA/AIS (n=17 PML, n=33 tumor samples), and GSE193725 which contained a mixture of paired normal/PML, normal/tumor, and independent samples from 42 total patients (n=23 PML, n=10 normal samples, n=18 tumor samples) (C) Heatmap of resulting matrix after summing binarized gene adjacency matrices from each of the 4 PML datasets and filtering to identify highly connected genes (n=1628 genes). Heatmap shades indicate number of datasets that the genes are correlated in, ranging from 0 (white) to 4 (deep red). Rows/columns are semi-supervised by modules identified using *igraph* clustering, with color annotations indicating each module (D) Heatmap of GSVA scores computed from each PML-derived gene module. Row labels indicate pathway enrichment of the nine modules, and row colors correspond to modules identified in Panel C. Columns are hierarchically clustered using *ward.d2* similarity of the Euclidean distance of samples (E) Boxplots of GSVA scores computed for each PML-derived module across the Lung PCA cohort samples (n=177 samples) stratified by module and colored by histology. Asterisk (*) indicates histology is significantly different from AAH samples (linear model with patient as a random effect, p<0.05)

**Table 1:**
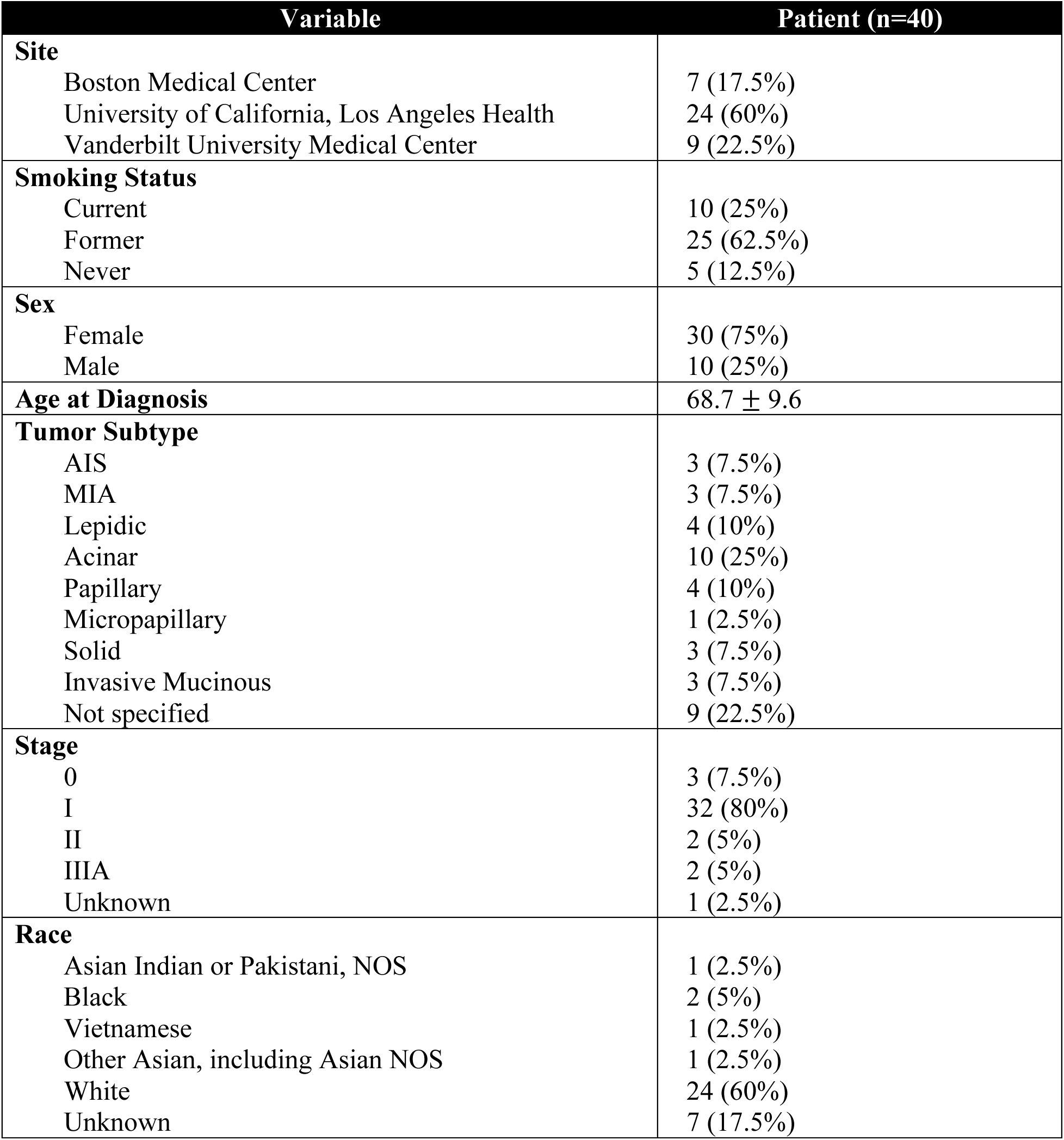
Clinical and Demographic Features of Lung PCA Cohort.

| Variable | Patient (n=40) |
| --- | --- |
| **Site** |  |
| Boston Medical Center | 7 (17.5%) |
| University of California, Los Angeles Health | 24 (60%) |
| Vanderbilt University Medical Center | 9 (22.5%) |
| **Smoking Status** |  |
| Current | 10 (25%) |
| Former | 25 (62.5%) |
| Never | 5 (12.5%) |
| **Sex** |  |
| Female | 30 (75%) |
| Male | 10 (25%) |
| **Age at Diagnosis** | 68.7 ± 9.6 |
| **Tumor Subtype** |  |
| AIS | 3 (7.5%) |
| MIA | 3 (7.5%) |
| Lepidic | 4 (10%) |
| Acinar | 10 (25%) |
| Papillary | 4 (10%) |
| Micropapillary | 1 (2.5%) |
| Solid | 3 (7.5%) |
| Invasive Mucinous | 3 (7.5%) |
| Not specified | 9 (22.5%) |
| **Stage** |  |
| 0 | 3 (7.5%) |
| I | 32 (80%) |
| II | 2 (5%) |
| IIIA | 2 (5%) |
| Unknown | 1 (2.5%) |
| **Race** |  |
| Asian Indian or Pakistani, NOS | 1 (2.5%) |
| Black | 2 (5%) |
| Vietnamese | 1 (2.5%) |
| Other Asian, including Asian NOS | 1 (2.5%) |
| White | 24 (60%) |
| Unknown | 7 (17.5%) |

We also used bulk RNA sequencing data from LUAD PML samples from three previously published datasets. Sivakumar et al. (GSE102511)^10^ performed RNA and targeted DNA sequencing to identify transcriptional changes and driver mutations from a cohort of East Asian patients diagnosed with predominantly stage I LUAD (RNA-seq samples included n=17 AAH from n=17 patients). This cohort is similar in study design to our cohort, profiling lesions adjacent to tumors, and provides increased patient diversity from a geo-cultural perspective. Yoo et al. (GSE166720)^15^ and Altorki et al. (GSE193725)^17^ performed RNA sequencing on patients who had only AIS/MIA (n=17 AIS/MIA from n=17 patients and n=23 AIS/MIA from n=23 patients, respectively) or stage I/II LUAD (**Figure 1B**). We selected these two studies to increase the number of AIS/MIA samples in our analysis and to include PMLs in patients without LUAD tumors because field cancerization^22^ may affect gene expression profiles of tissue adjacent to tumors.

### Identification of gene co-expression modules in LUAD premalignancy

To identify novel patterns of transcriptional heterogeneity among LUAD PMLs, we used a discovery-based approach to identify PML archetypes based on patterns of gene module co-expression. To accomplish this, we first derived a set of gene modules that consisted of genes with robust co-expression across multiple LUAD PML data sets (**Figure 1B**). To identify recurring gene co-expression relationships, we identified gene-gene pairs with strong expression correlation in at least 3 datasets, and further selected genes that demonstrated connections to multiple other genes, resulting in a total of 1628 genes (**Figure 1C**). The filtered gene connection matrix was clustered using *igraph*^23^ and 9 gene modules were identified (average number of genes=180, minimum=5, maximum=523). We observed, for each module and dataset, that genes within a module were significantly more correlated to the module gene set variation analysis^24^ (**GSVA**) score than to genes outside the module (**Supplementary** Figure 1A, Kolmogorov–Smirnov test, p<0.0001), confirming consistent co-expression of the modules across data sets. Modules genes were significantly enriched (FDR<0.05) for “Biological Process” Gene Ontology terms via *EnrichR*^25^ and included diverse processes such as extracellular matrix organization, cell cycle, and cytokine signaling. Module GSVA scores across all samples were orthogonal (average module-module GSVA score Pearson R=0.12 +/- 0.097) in accordance with their unique biology. Together, these modules did not separate samples into clusters that were enriched for dataset, sex, or smoking status (chi-square using *k*=2 clusters as shown in first split of dendrogram in **Figure 1D**, p=0.35, p=1, p=0.062, respectively). To quantify the relationship of individual module GSVA scores with histology, using AAH samples as the basis for comparison (**Table 2)**, we found that the cell cycle and extracellular matrix organization modules were higher in tumors compared to AAH, with cell cycle also being higher in AIS/MIA compared to AAH. In contrast, the regulation of endothelial proliferation module was increased in normal samples compared to AAH (**Figure 1E**, linear model with patient as random effect). These associations were also preserved in the 3 additional datasets used to derive the modules (**Supplementary** Figure 1B**, C, and D**). Across datasets, the gene modules had unique conserved patterns of co-expression and biology, and demonstrate significant associations with histology.

**Table 2:** Comparison of GSVA scores for PML-derived modules among Lung PCA samples.

| Direction of GSVA scores compared to AAH | Histology comparison | Modules |
| --- | --- | --- |
| Increased (p<0.05) | ADC | • Cell cycle<br>• Extracellular matrix organization<br>• T-cell activation<br>• Inflammatory response |
|  | AIS/MIA | • Cell cycle<br>• Inflammatory response |
|  | Normal | • Regulation of endothelial cell proliferation |
| Decreased (p<0.05) | ADC | • Regulation of endothelial cell proliferation |
|  | AIS/MIA | • Cytokine signaling<br>• Regulation of endothelial cell proliferation |
|  | Normal | • None |

### Gene co-expression modules identify four distinct archetypes of PMLs

We hypothesized that the module co-expression heterogeneity among PMLs might be driven by different molecular phenotypes. We therefore sought to use the PML-derived gene modules to define archetypes^19,26^ of LUAD PMLs, an approach previously used to identify phenotypic heterogeneity in small cell lung cancer^21^. To avoid biasing the results towards modules with larger numbers of genes, we used the GSVA scores for each module (**Figure 1D**) as the input and included all PML samples used to identify the gene modules. We determined that four archetypes best fit the data based on a graph of explained variance versus *k* (**Supplementary** Figure 2A, *Methods*) and a t-ratio^27^ permutation test (**Supplementary** Figure 2B, *Methods*). Samples were scored based on their distance from each archetype (**Figure 2A**) and the octile of samples (12.5%), with the highest similarity scores were categorized as belonging to an archetype (**Figure 2B**). If samples had overlapping primary labels (n=6 samples, 4.4% of samples) or did not have a primary label from any of the archetypes (n=75, 55% of samples), we labeled them as “secondary”. We labeled the four archetypes as: normal-like (**A1, green**, n=12 samples, mean distance from A1 center=-0.99 +/-0.15) inflammation (**A2, blue**, n=13 samples, mean distance from A2 center=- 0.99 +/-0.10), cell adhesion (**A3, purple**, n=16 samples, mean distance from A3 center=-0.98 +/- 0.16), and proliferation (**A4, yellow**, n=15 samples, mean distance from A4 center=-0.93 +/-0.19). None of the four archetypes have categorically designated samples that are significantly further or closer to the archetype center compared to each of the other archetypes (Wilcox test, p>0.4 for all comparisons).

**Figure 2:**
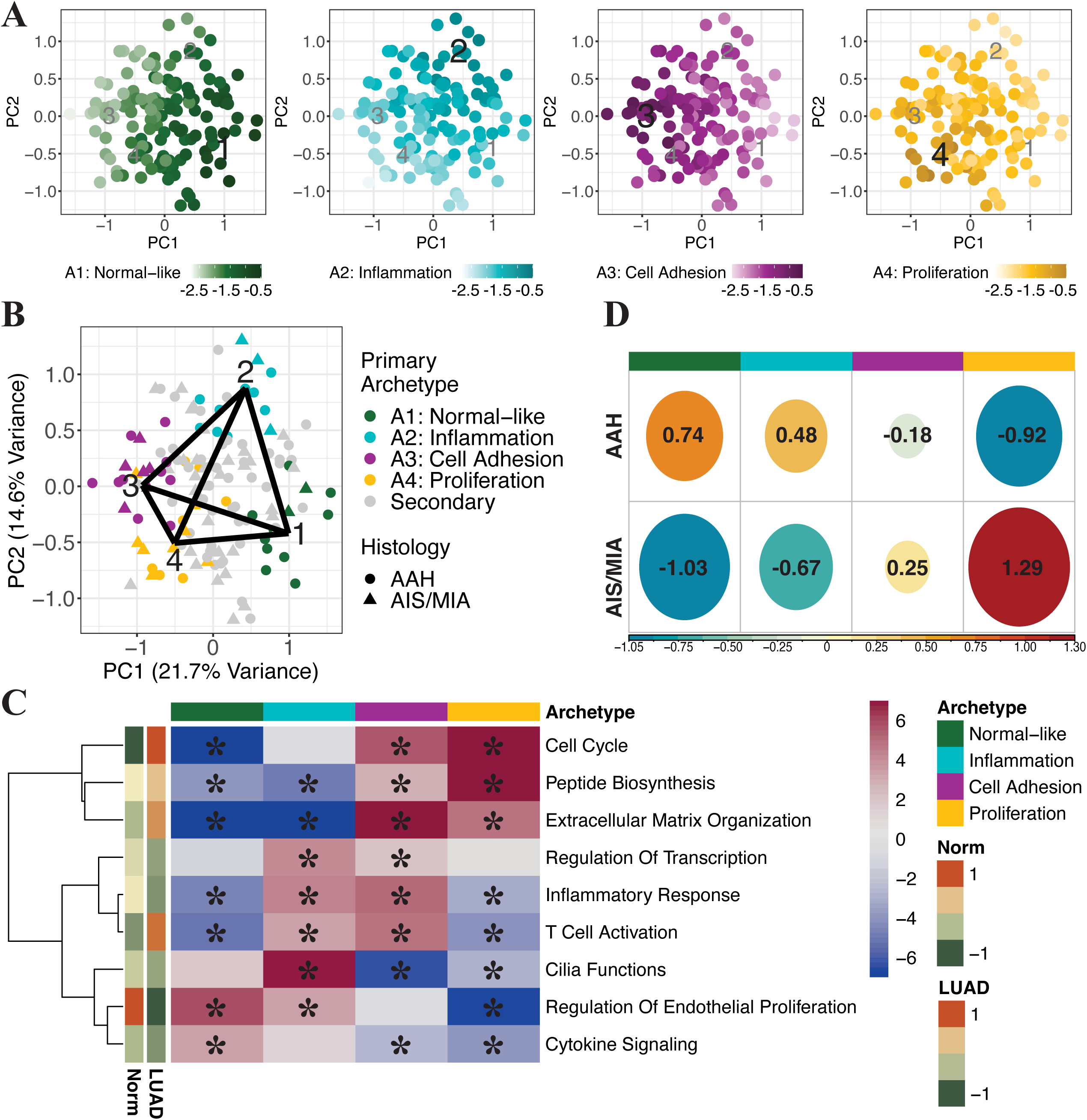
Archetype analysis on PMLs demonstrates correlation of archetypes with molecular and clinical phenotypes. (A) Scatterplots of first two principal components computed from GSVA scores for PML-derived modules across PML samples from all four cohorts. Each plot is colored by the distance of each sample from one archetype center. Each sample has four associated distances—one for each archetype vertex. More negative numbers reflect samples further away from the vertex (B) Scatterplot of first two principal components computed from GSVA scores for PML-derived modules for all PML samples. Samples in the top 0.125 proportion of distances from each archetype were designated as primary archetypes, while other samples were labeled as “secondary”, with colors representing samples closest to each respective archetype. Shapes indicate histology of samples (C) Heatmap of t-statistics from linear models of GSVA scores for each PML-derived module (rows) modeled as a function of distance from each archetype (columns). Asterisk (*) indicates FDR<0.05 (linear model adjusted for study with random effect for patient). Rows are hierarchically clustered using *ward.d2* similarity of the Euclidean distance. Row annotations represent the median GSVA score for each module in Lung PCA samples of either normal tissue or tumor (D) Visualization of chi-square residuals from chi-square test of association between archetypes and histology. Plot shows contribution of each comparison in the matrix to the test statistic, where circle size is proportional to contribution. Positive (red) values specify a positive association between the corresponding row and column variables (chi-square test, p=0.18, n=37 AAH, n=19 AIS/MIA)

Next, we characterized the archetypes using the premise that samples close to an archetype share specific biological functions^19,26^. We identified PML-derived gene module scores that were significantly associated with distance from each archetype (**Figure 2C**, t-statistic from linear model adjusted for study, with patient as random effect, FDR<0.05 indicated with asterisk), and Molecular Signatures Database “Hallmark” gene signatures^28,29^ that were significantly enriched among genes correlated with archetype distances using Gene Set Enrichment Analysis^30,31^ (**GSEA**) (**Supplementary** Figure 3, only pathways with positive normalized enrichment score and FDR<0.05 are shown). Samples close to the normal-like archetype were positively associated with increased GSVA scores for the regulation of endothelial cell proliferation module (FDR<0.0001), which was also significantly higher in samples with normal histology compared to AAH lesions (**Figure 1E**, p=0.0011). The normal-like archetype was also associated with pathways that may regulate cell proliferation (P53) or activate apoptotic pathways (UV response up). Samples closer to both the inflammation and cell adhesion archetypes had significantly higher GSVA scores for the T-cell activation (FDR<0.0001) and inflammatory response (FDR<0.0001) modules, and up- regulation of Hallmark immune-related pathways (**Supplementary** Figure 3). The extracellular matrix organization module expression was a marked difference between these archetypes as samples close to the cell adhesion and inflammation archetypes were positively and negatively associated (FDR<0.0001) with the module, respectively. Samples close to the cell adhesion archetype also had increased expression of Hallmark angiogenesis, epithelial-to-mesenchymal transition, and *Wnt* and hedgehog signaling pathways, suggesting a molecular phenotype linked to cell migration. Samples close to the proliferation archetype were most positively associated with increased GSVA scores for the cell cycle module (FDR<0.0001), which was also significantly higher in tumor samples compared to AAH samples (**Figure 1E**, p<0.0001). Proliferation archetype samples also had increased expression of genes in pathways associated with cell proliferation, metabolic functions, and oxidative stress in the Hallmark gene set.

We also determined the associations between the archetypes and histology. The proliferation archetype was the only archetype significantly associated with AIS/MIA versus AAH histology (logistic regression adjusted for dataset with patient modeled as random effect, n=85 AAH and n=52 AIS/MIA, p=0.072 for normal-like, p=0.062 for inflammation, p=0.86 for cell adhesion, p=0.021 for proliferation). Among samples with a categorical archetype designation, AIS/MIA samples were positively enriched, but not statistically significantly, in the proliferation archetype samples compared to other archetypes (**Figure 2D**, **Supplementary Table 1**, chi-square test, p=0.18, n=37 AAH, n=19 AIS/MIA). There was no association of the archetypes with study (**Supplementary** Figure 4A, chi-square, p=0.33), smoking status (**Supplementary** Figure 4B, chi-square, p=0.29), or sex (**Supplementary** Figure 4C, chi-square, p=0.65). Overall, the archetypes had distinct and strong associations with biological pathways but were not strongly associated with severity of PML histology.

### Archetypes are associated with the somatic mutation landscape in premalignancy

To identify the relationship between LUAD PML archetypes and somatic mutations, we first examined mutant allele tumor heterogeneity (**MATH**)^32^ across samples in the Lung PCA cohort with exome sequencing data (n=26 LUAD, n=12 AIS/MIA, n=52 AAH samples from n=25 patients). MATH is a measure of the mutant allele fraction, where heterogenous tumors with many genetically distinct subclones will have a higher MATH score. Tumor samples had a significantly higher MATH score compared to PMLs (linear model with random effect for patient, p=0.00012). Next, we found significant positive enrichment of driver mutations in tumor samples compared to AAH and AIS/MIA samples for both *EGFR* (**Supplementary Table 2**, chi-square, p=0.018) and *TP53* (**Supplementary Table 3**, chi-square, p=0.0024) mutations. There was no association with driver mutation presence and histology for *KRAS* mutations (**Supplementary Table 4**, chi-square, p=0.99). For patients with available exome sequencing data for both tumor and PMLs (n=69 PMLs and n=21 LUAD from n=15 patients), we computed the Jaccard similarity based on shared deletions, insertions, and single nucleotide polymorphisms to determine relatedness between individual PMLs and their respective paired tumor. The mean similarities were higher between tumors and PMLs from the same patient (0.0050 +/- 0.017, maximum=0.11) compared to the mean similarities between tumors and PMLs from unrelated patients (0.00012+/-0.00017, maximum=0.00063) (Wilcoxon rank sum test, p=0.020). The similarity between PMLs and tumors from the same patient was overall low, however, indicating few shared somatic alterations. These results indicate that tumors have increased genomic instability compared to PMLs, as previously observed^10^.

We next assessed the relationship between archetypes and the PML somatic mutational landscape. Lung PCA PMLs assigned to the proliferation archetype had a significantly increased MATH score (**Figure 3A**, linear model with random effect for patient, p<0.0001) compared to all other samples. When comparing the MATH score of the tumor samples to the categorical archetype samples, the normal-like, inflammation, and cell adhesion archetypes had lower MATH scores (p=0.16, p=0.0022, and p=0.032, respectively). There was no significant difference between tumor and categorical proliferation archetype PML sample MATH scores (p=0.76), indicating a similar degree of MATH in proliferation samples and LUAD samples. The mean Jaccard distance for PMLs assigned to the proliferation, cell adhesion, inflammation, and normal-like archetypes and their respective tumor samples was 0.014 (n=5 patients, maximum=0.11), 0.00016 (n=3 patients, maximum 0.0012), 0.00025 (n=3 patients, maximum 0.0017), and 0.0021 (n=3 patients, maximum 0.0033), respectively. Samples assigned to the proliferation archetype were the most like their paired tumor samples, but this does not meet statistical significance compared to the normal-like, inflammation, or cell adhesion archetypes (Wilcoxon rank sum test comparing other archetypes to proliferation, p=0.26, p=0.090, and p=0.061, respectively). We also examined driver mutations in these PMLs and GSE102511 samples (n=13 AAH) with available targeted DNA sequencing data, focusing on mutations that appeared in at least 4 PMLs. We found that archetypes have differential enrichment of driver mutations, (**Supplementary Tables 5-8**). Samples closer to the proliferation archetype were more likely to be mutated for *EGFR* (p=0.0021) or *JAK3* (p=0.062) (**Figure 3B**, logistic regression adjusted for study and with random effect for patient, asterisk indicates p<0.05). These results suggest that PMLs have fewer somatic alterations and a weak genetic relationship with their paired tumors. However, the proliferation archetype PMLs were positively enriched for *EGFR* mutations, similar to tumors, and tended to be more similar to tumors compared to other archetypes.

**Figure 3:**
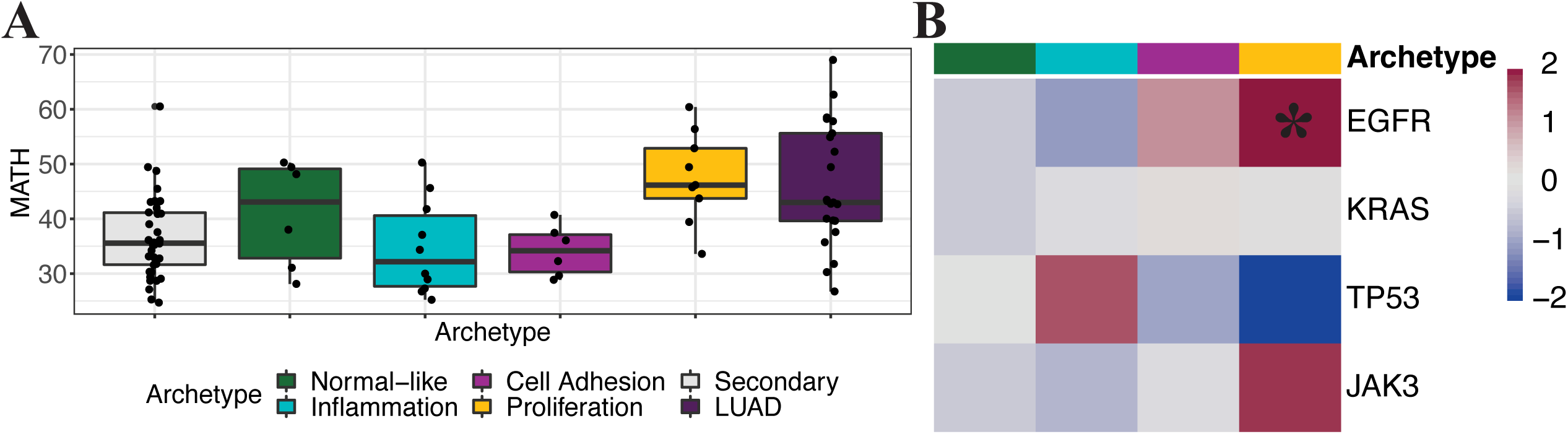
Archetypes are associated with the somatic mutation landscape. (A) Boxplot of mutant tumor allele heterogeneity score for Lung PCA PML samples with categorical archetype designation (n=6 normal-like, n=10 inflammation, n=6 cell adhesion, n=9 proliferation, and n=38 secondary) and Lung PCA tumor samples (tumor samples were not included in archetype analysis, n=21 samples) (B) Heatmap of z-statistic for absence/presence of each driver mutation (rows) modeled as a function of distance from archetype (columns) in Lung PCA (n=57 AAH and n=12 AIS/MIA samples) and GSE102511 (n=13 AAH samples) samples with available exome sequencing data. Asterisk (*) indicates p<0.05 (logistic regression adjusted for study with patient as a random effect)

### Archetypes are associated with altered immune phenotypes

To further understand the relationship of the archetypes to altered immune microenvironments, we used signatures from the study by Sivakumar et al. (GSE102511)^10^, which defined immune-specific gene clusters differentially expressed between normal, AAH, and LUAD samples (**Figure 4A**), and cell type-specific signatures from Bischoff et al.^33^ derived from single-cell RNA sequencing of LUAD (**Figure 4B**). For each signature, we computed GSVA scores and examined if these scores varied based on distance from each archetype (linear model adjusted for study and with random effect for patient, asterisk indicates FDR<0.05). On both an immune pathway level (Sivakumar et al., **Figure 4A**) and a cell type level (Bischoff et al., **Figure 4B**), samples close to the cell adhesion and proliferation archetypes were significantly enriched for both pro-tumor B-cell signaling and immune checkpoint pathways, and cancer-associated myofibroblasts and exhausted CD8^+^ T-cells associated with a poor prognosis. Conversely, samples close to the normal-like and inflammation archetypes showed the opposite pattern of expression. The similarities and differences between the archetypes were also found when looking at non-immune focused gene co-expression clusters identified by Sivakumar et al. (**Supplementary** Figure 4D). Despite the noted similarity between the proliferation and cell adhesion archetypes, samples closer to the proliferation archetype also had a significant decrease in both anti-tumor immune responses and signatures for natural killer and T-cells associated with better prognosis.

**Figure 4:**
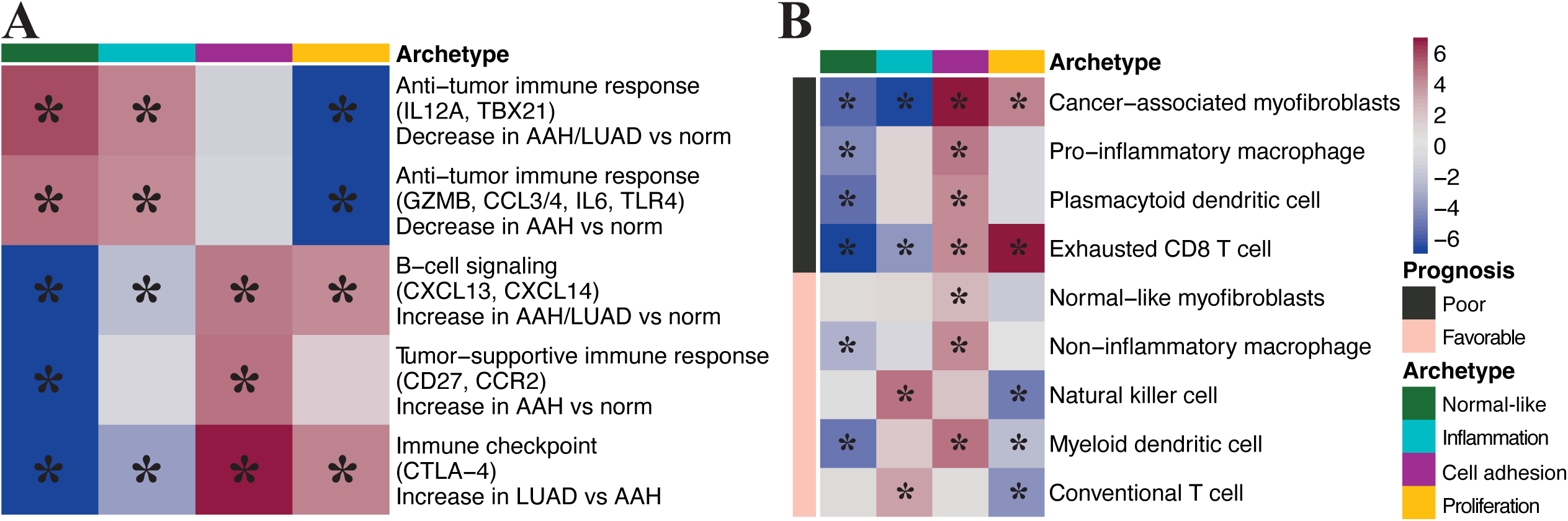
Archetypes are associated with signatures of LUAD aggression and prognosis. (A) Heatmap of t-statistics for linear model of GSVA scores for each Sivakumar et al. gene signatures that differ between normal, AAH, and tumor samples (rows) modeled as a function of distance from each archetype (columns). Asterisk (*) indicates FDR<0.05 (linear model adjusted for study with random effect for patient) (B) Heatmap of t-statistics for linear model of GSVA scores for Bischoff et al. cell type gene signatures (rows) modeled as a function of distance from each archetype (columns). Asterisk indicates FDR<0.05 (linear model adjusted for study with random effect for patient)

Using multiplex immunofluorescence (**MIF**) performed on a subset of Lung PCA samples as described by Yanagawa et al.^34^ (**Supplementary** Figure 4E, n=39 samples total, n=30 AAH, n=9 AIS/MIA), we examined changes in the immune environment between archetypes, modeling cell density as a function of distance from each archetype. Samples closer to the proliferation archetype demonstrated a decrease in natural killer T-cells (p=0.018), and a modest decrease in CD4^+^ (p=0.16) and CD8^+^ (p=0.26) T-cell density. The added resolution of T-cell type signatures identified by Bischoff et al. (**Figure 4B**) showed a statistically significant increase in exhausted CD8^+^ T-cells and a decrease in conventional T-cells, suggesting that specific CD4^+^and CD8^+^ T-cell populations may be important to the proliferation archetype. The reverse pattern is observed for samples closer to the inflammation archetype, which have a significantly increased CD8^+^ T-cell (p=0.0066) and modestly increased CD4^+^ T-cell (p=0.28) and natural killer T-cell (p=0.28) density in the MIF analysis, and significantly increased natural killer T-cell and conventional T-cell density in the Bischoff et al. signatures. These results aid in the characterization of the archetype immune microenvironments, where the archetypes likely represent a spectrum of progressive changes to immune pathways and cell types.

### Archetypes are related to tumor histopathologic and clinical features in multiple studies of LUAD

To better understand the relationship of the archetypes to features of LUAD including stage, histopathologic features, and prognosis, we fit data from multiple LUAD datasets to the PML polytope. We utilized transcriptional data from LUAD tumors in the Lung PCA cohort (n=49 samples from n=30 patients), the Yoo et al study (GSE166720^15^, n=33 samples, n=33 patients), a prospective multi-center study of stage I, II, and III lung cancer in the United Kingdom (**TRACERx**^35^, n=460 samples, n=186 patients), the Director’s Challenge Consortium for the Molecular Classification of stage I, II, and III LUAD (GSE68465^36^, n=362 samples, n=362 patients), stage I and II LUAD from studies by Okayama et al. and Yamauchi et al. (GSE31210^37,38^, n=226 samples, n=226 patients), and stage I LUAD from Rousseaux et al. (GSE30219^39^, n=85 samples, n=85 patients). Prior to fitting each dataset to the PML-derived polytope using module GSVA scores, we confirmed that there was a stronger correlation between genes within versus outside each module to the module GSVA scores in independent LUAD datasets (**Supplementary** Figure 5, Kolmogorov–Smirnov test, p<0.0005 for every module and every dataset). After the LUAD datasets were fit to the PML polytope (**Supplementary** Figure 6), we observed similar distances between categorical archetype samples and their respective archetype centers for each dataset (**Supplementary** Figure 7), and a consistent relationship between sample distance from each of the archetypes and the module GSVA scores (**Supplementary** Figure 8). Additionally, in the Lung PCA data (n=49 LUAD, n=48 normal, n=80 PML), the inclusion of the LUAD samples only changed the categorical archetypes of assignment of 2 PML samples (2 cell adhesion archetype PML re-assigned to “secondary”) (**Figure 5A**). These results indicated that PML- derived module co-expression patterns and their relationships to the four archetypes were preserved across LUAD datasets.

**Figure 5:**
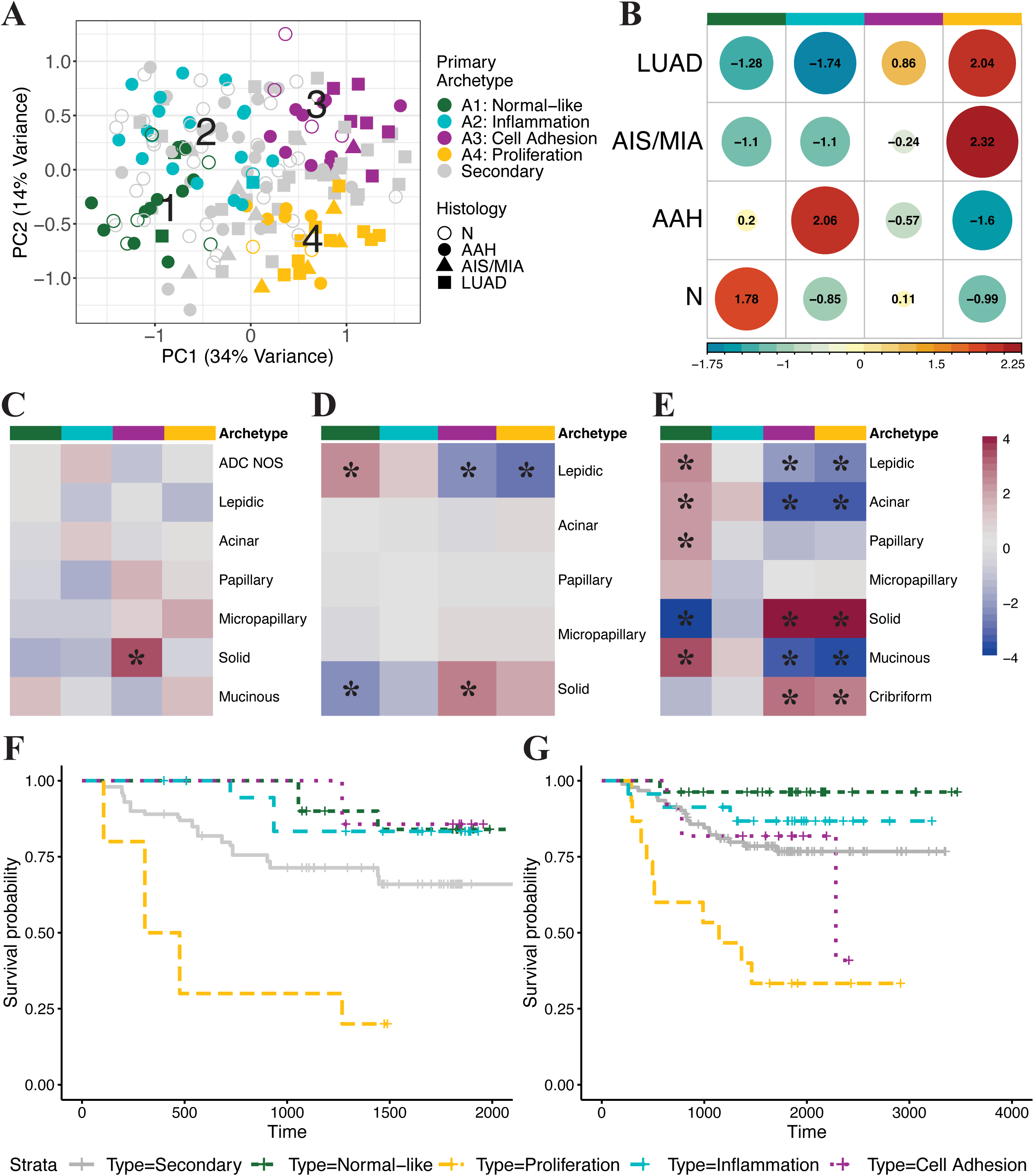
Projection of archetypes into cancer data stratifies cohorts by lesion features and prognosis. (A) Scatterplot of first two principal components computed from GSVA scores for PML-derived modules in full Lung PCA cohort, with annotated projection of PML archetypes into Lung PCA principal component space. Samples in the top 0.125 proportion of distances from each archetype were designated as primary archetypes, while other samples were labeled as “secondary”, with colors representing samples closest to each respective archetype. Shapes designate sample histology (B) Visualization of chi-square residuals from chi-square test of association of archetypes with histology in Lung PCA samples. Plot shows contribution of each cell to the test statistic, where circle size is proportional to cell contribution. Positive (red) values specify a positive association between the corresponding row and column variables (p=0.00053) (C-E) Heatmap of t-statistics for linear model of distance from each archetype (columns) modeled as a function of histologic tumor type (rows) for: (C) Lung PCA tumor samples (D) GSE166720 tumor samples and (E) TRACERx tumor samples, with histology indicating overall histology of tumor. Asterisk (*) indicates significant relationship of archetype to histology (F-G) Cox Proportional-Hazard Model of disease-free survival probability for samples closest to each archetype in (F) TRACERx stage I tumor samples (n=158 samples, n=77, patients, p=0.00016, Hazard Ratio=4.50 for samples assigned to proliferation archetype) (G) GSE31210 stage I tumor samples (n=168 samples, n=168 patients, p=0.00027, Hazard Ratio=4.12 for samples assigned to proliferation archetype)

In the Lung PCA dataset, inclusion of LUAD samples led to a significant association between the sample archetypes and histology (**Figure 5B**, **Supplementary Table 9**, chi-square, p=0.00053). The proliferation archetype was enriched with LUAD and AIS/MIA samples, the inflammation archetype with AAH samples, and the normal-like archetype with normal samples. Next, we examined the archetype relationships between PMLs and their paired tumors. We observed that paired samples demonstrated increased similarity (Wilcox test, p=0.0021) in the distance from archetype centers of PMLs and their paired tumors (n=58 PML, n=26 LUAD, n=21 patients; cosine similarity, mean=0.96+/-0.044) compared to PMLs and unrelated tumors (cosine similarity, mean=0.95+/-0.043). The similarity is positively but not significantly correlated with the Jaccard similarity computed with the exome sequencing for paired tumors and PMLs (Pearson R=0.15, p=0.28). To understand if categorical PML archetype influences tumor-PML pair similarity, we computed the cosine similarity between PMLs assigned to either the normal-like (n=8 PML, n=7 LUAD, n=6 patients) or proliferation (n=7 PML, n=6 LUAD, n=5 patients) archetypes with paired tumors (**Supplementary** Figure 9). The normal-like PMLs had a significantly lower similarity (mean=0.92+/-0.062) than PMLs assigned to the proliferation archetype (mean=0.97+/-0.025) from their paired tumors (Wilcox test, p=0.00017). There is no significant difference between the similarity of proliferation archetype PMLs and paired tumors compared to the cell adhesion (Wilcox test, p=0.88) or inflammation archetype (Wilcox test, p=0.95) PMLs and paired tumors. These results indicate that paired PMLs and tumors have relatively weak similarity, similar to the findings from the exome sequencing data, and that normal-like archetype PMLs are the least similar in archetype space to their paired tumor samples.

Next, we examined the relationship between the PML-derived archetypes and LUAD stage, histopathologic features and lymph node metastasis. Samples closer to the proliferation archetype were associated with higher stage (stage II and above) and samples closer to the normal-like archetype were associated with stage I samples, respectively, in TRACERx (p=0.020, p=0.071), GSE68465 (p=0.00013, p=0.0042), GSE31210 (p<0.0001, p<0.0001), and Lung PCA LUAD samples(p=0.031, p=0.010). Using the overall predominant histology of patient tumors (as opposed to individual sample histology), we found that samples closer to the cell adhesion archetype were enriched with the aggressive solid histologic pattern in Lung PCA (**Figure 5C**, linear model with patient as a random effect and adjusted for stage, p=0.0010), Yoo et al. (**Figure 5D**, linear model, p=0.010), and TRACERx LUAD samples (**Figure 5E**, linear model with patient as a random effect and adjusted for stage, p<0.0001). In TRACERx, samples from patients with solid predominant tumors were also significantly closer to the proliferation archetype (p<0.0001). Samples from tumors with the less aggressive lepidic pattern were significantly closer to the normal-like archetype in both the Yoo et al. and TRACERx LUAD samples (p=0.027, p=0.015, respectively). In a subset of Lung PCA LUAD samples collected from Boston Medical Center (n=6 samples from n=6 patients), the invasive non-lepidic portion of the tumor was measured and found to be positively correlated with closeness to the proliferation archetype and negatively correlated to the normal-like archetypes (r=0.77, p=0.076 and r=-0.78, p=0.067, respectively). In TRACERx, the size of the non-lepidic portion of the tumor was positively correlated with closeness to the proliferation archetype (r= 0.30, p<0.0001). In categorically assigned GSE68465 samples (n=210 samples, n=210 patients), archetypes were differentially associated with tumor grade (**Supplementary** Figure 10A, **Supplementary Table 10**, chi-square, p<0.0001) and lymph node metastasis (**Supplementary** Figure 10B, **Supplementary Table 11**, chi-square, p=0.0038).

Samples assigned to the normal-like archetype were positively associated with low grade tumors, while samples assigned to the proliferation archetype were positively associated with high grade tumors and lymph node metastasis. These findings were consistent when examining the relationship between sample distance from the proliferation archetype and tumor grade or lymph node metastasis in the full GSE68465 dataset (p=0.0039, p=0.0036, respectively), in Lung PCA tumor samples (linear model with patient as random effect, p=0.0051, p=0.081), and TRACERx (linear model with patient as random effect, tumor grade not available, p=0.031 for lymph node metastasis). In the TRACERx dataset where the presence of lymphovascular invasion was annotated, samples closer to the normal-like archetype were associated with the absence of lymphovascular invasion while the opposite was true for the proliferation archetype (linear model with patient as random effect, p=0.020, p=0.021, respectively). Taken together, the proliferation archetype was associated with the aggressive solid predominant histologic pattern, tumor invasive size, high tumor grade, lymph node metastasis, and lymphovascular invasion suggesting that this archetype is associated with aggressive LUAD compared to the other archetypes.

Given the association between the proliferation archetype and aggressive LUAD features, we wanted to determine if the proliferation archetype was associated with poor disease-free survival. As stage I LUAD are likely to be treated with surgical resection without additional therapy, we choose to focus the analysis on this subset of LUAD. In both the TRACERx stage I data (n=158 samples, n=77 patients, **Figure 5F**) and GSE31210 stage I data (n=168 samples, n=168 patients, **Figure 5G**), there was a significant difference in disease-free survival time between samples assigned to the different categorical archetypes, with shorter disease-free survival in samples assigned to the proliferation archetype (p=0.00016, Hazard Ratio=4.50 and p=0.00027, Hazard Ratio=4.12, respectively). Using sample distance to the proliferative archetype, we found that stage I TRACERx LUAD samples closer to the proliferation archetype trended towards shorter disease-free survival (p=0.14, Hazard Ratio=3.31). Stage I LUAD samples from GSE31210 and GSE30219 closer to the proliferation archetype were also significantly associated with worse prognosis (p=0.00086, Hazard Ratio=3.31 and p=0.0088, Hazard Ratio=3.08, respectively). These results suggest that the proliferation archetype is associated with worse disease-free survival in stage I LUAD samples in multiple independent datasets.

## DISCUSSION

Wider adoption of lung cancer screening with low-dose CT has increased the detection of pure ground glass opacities (**GGO**) that can represent premalignant histology^40^, presenting a need to develop biomarkers that can distinguish between indolent and aggressive PMLs that will progress to invasive lung cancer^6^. Prior work profiling LUAD PMLs using bulk RNA and exome sequencing focused on identifying similarities and differences between PMLs and tumors^8–15,17^. However, a recent study of East Asian never smokers found few differentially expressed genes by pathologic stage, but more changes associated with radiologic features^18^. There is an unmet need to characterize molecular heterogeneity in LUAD PMLs and find subtypes associated with prognosis^41,42^ to develop biomarkers to prevent overtreatment of indolent PMLs and hasten treatment of aggressive PMLs, such as those detected via current lung cancer screening programs^7,43^. To this end, using LUAD PMLs profiled by RNA-seq from four cohorts, we identified nine conserved gene co-expression modules. Of these modules, four had significantly altered expression between pathologic stage (AAH compared to AIS/MIA) in the Lung PCA cohort. These nine modules were subsequently used to define four archetypes—normal-like, inflammation, cell-adhesion, and proliferation—that represent different gene expression states among PMLs, and are associated with activation of genes involved in distinct biological pathways, immune microenvironments, and somatic mutation profiles. The gene expression patterns that define each archetype were able to discriminate between aggressive and indolent LUAD across multiple studies, linking PML molecular heterogeneity and prognostic histopathological features in LUAD.

PMLs representative of the proliferation archetype were associated with the presence of *EGFR* driver mutations, increased mutational heterogeneity, an altered immune landscape, and increased expression of cell proliferation and metabolic pathways, suggestive of rapid cell growth. Previously, the frequency of *EGFR* mutations had been found to increase as lesions progress along the histologic spectrum from AAH to AIS/MIA to LUAD^18^, and are more frequent in tumors compared to AAH^10^. Our study similarly found that *EGFR* mutations were enriched in tumor samples compared to PMLs. However, within PMLs, we found the proliferation archetype (enriched for AIS/MIA) was enriched for *EGFR* mutation compared to PMLs from all other archetypes. We also demonstrated that proliferation archetype PMLs have the highest levels of intra-lesion mutational heterogeneity, similar to tumor samples. The proliferation archetype was also associated with both tumor-supportive immune pathways identified by Sivakumar et al.^10^ and signatures for pro-inflammatory macrophages and exhausted CD8^+^ T-cell populations identified by Bischoff et al. as being associated with a poor prognosis^33^. A spatial analysis of AAH, AIS/MIA and LUAD similarly found that regions with decreased innate inflammatory and increased immune evasion responses were associated with increases in PDL1^+^Ki67^+^ epithelial cells^44^, which is consistent with the enrichment of the proliferation archetype for cell cycle-related gene expression. Interestingly, tumors most similar to the proliferation archetype had significantly decreased disease-free survival time, were enriched for aggressive histologic patterns and invasive features, and had higher grade compared to tumors most similar to the other archetypes. Collectively, these findings indicate that some PMLs share gene expression patterns with aggressive LUAD. This further suggests that patients with PMLs similar to the proliferation archetype may benefit from more frequent CT screening, targeted interception strategies, or resection.

In contrast to the proliferation archetype, PMLs representative of the normal-like archetype shared module gene expression patterns with histologically normal tissues adjacent to tumors, or with lepidic tumors in studies of LUAD, and lacked significant associations with known driver mutations. Normal-like archetype samples had increased expression of genes involved in pathways regulating endothelial cell proliferation, potentially reflecting the critical role endothelial cells have in alveolar gas exchange in normal lung tissue^44^. Additionally, normal-like archetype samples showed increased expression of genes in anti-tumor immune pathways^10^ and decreased expression of tumor-associated cell type signatures such as cancer-associated myofibroblasts, pro-inflammatory macrophages, and exhausted CD8^+^ T-cells^33^. This potentially represents a subset of lesions similar to those observed by Krysan et al. that exhibit active immune surveillance^12^. Normal-like archetype PMLs may represent a subset of lesions that are unlikely to progress to cancer^7^ and could be managed conservatively via repeat CT imaging.

The inflammation and cell adhesion archetypes share module gene expression with both the normal-like and proliferation archetypes and potentially represent intermediate LUAD PML phenotypes. The inflammation archetype, similar to the normal-like archetype, demonstrated elevated expression of anti-tumor immune pathways^10^, and expression of cell type-specific markers of natural killer and conventional T-cells that were associated with a more favorable prognosis^33^. Collectively, these data suggest that inflammatory archetype PMLs may be recognized by the immune system^10–12,14,15,34^. Samples representative of the cell adhesion archetype shared some features with the proliferation archetype, including increased expression of immune checkpoint pathways^10^ and signatures for cancer-associated myofibroblasts and exhausted CD8^+^ T-cells associated with a less favorable prognosis^33^. In contrast with proliferation archetype samples, lesions close to the inflammation and cell adhesion archetypes had significantly lower intra-lesion mutational heterogeneity compared to tumors. Unlike the proliferation archetype, LUAD tumors closest to the cell adhesion archetype were not associated with decreased disease-free survival, higher stage, higher grade, or lymphovascular invasion. The cell adhesion archetype samples displayed increased expression of the PML-derived extracellular matrix module, the Hallmark epithelial-to-mesenchymal transition pathway, and higher expression of a cancer-associated myofibroblast gene signature^33^. These findings not only suggest that PMLs vary with respect to their level of stromal interaction, but also raise the question about the role of these stromal interactions in early tumorigenesis, as recently highlighted in studies of AIS/MIA^17,18^. Our characterization suggests the archetypes may represent a spectrum of indolent to aggressive transcriptional patterns among LUAD PMLs, with the cell adhesion archetype representing an intermediate between the inflammation and proliferation archetypes. The alterations in the stromal and immune features in the cell adhesion archetype compared to the proliferation archetype may modulate unchecked proliferation, leading to a less aggressive phenotype.

To determine if the transcriptional patterns that define the PML archetypes are associated with progression to lung cancer, future multi-omics studies are needed to molecularly, radiologically and pathologically phenotype PMLs over time. Like many published works characterizing LUAD PMLs, the lesions profiled in this study were identified in the margins of resected tumors and may be impacted by field cancerization^22^. LUAD PMLs are small and located deep in lung parenchyma, making them inaccessible to biopsy by standard bronchoscopy, which complicates longitudinal sampling^42^. As techniques such as robotic-assisted^45^ and electromagnetic navigation bronchoscopy^46^ become more widely available and accurate, and evidence continues to build about the diverse natural history of LUAD PMLs^7,40,41^, sampling of PMLs, pure GGOs, and part-solid GGOs may become more routine both clinically and to study disease.

In summary, our archetyping approach identified PML-variable features across multiple datasets to characterize PML heterogeneity. These features associate with pathway dysregulation and LUAD tumor stage, histopathologic features, and prognosis. Molecular signatures that can distinguish between indolent and aggressive PMLs may help personalize lung cancer screening protocols, prevent overdiagnosis, and allow for treatment that intercepts the development of aggressive LUAD via surgery and/or therapy to prevent cancer progression. Future longitudinal studies that monitor and sample nodules over time will be necessary to determine the direct relationship between PML archetypes, progression to lung cancer, and the histopathologic and prognostic features of the LUAD that may develop from a PML.

## METHODS

### Study population and sample collection

Patients that underwent resection for LUAD consented to the use of remanent tissue for research at University of California, Los Angeles (n=24 patients, n=119 normal, PML, and tumor samples post-filtering), Vanderbilt University Medical Center (n=9 patients, n=30 normal, PML, and tumor samples post-filtering), and Boston Medical Center (n=7 patients, n=28 normal and tumor samples post-filtering). Cases were reviewed to identify tumors where adjacent tissue contained premalignant histology as part of the Lung Pre-Cancer Atlas supported by the National Cancer Institute Consortium for Integrated Molecular, Cellular, and Imaging Characterization of Screen-Detected Lung Cancer and the Human Tumor Atlas Network. Tissue areas from all resected formalin-fixed paraffin-embedded (**FFPE**) tissues blocks for each case were selected following H&E review by a pathologist at each center. Samples selected for this study had normal, tumor, and premalignant histology confirmed by two independent pathologists, and any samples without agreement were removed from the analysis (n=26 samples removed).

Contamination of PML samples with other tissue may lead to the dilution and possible loss of low abundance molecular signals from premalignant cells that are necessary for understanding early cancer pathogenesis. Therefore, we utilized laser capture micro-dissection (**LCM**) to isolate normal, PML, and tumor samples from FFPE tissue samples, and to enrich the study samples with the cells of interest and minimize contamination with the surrounding tissue. RNA and DNA were isolated from the LCM tissue and profiled with bulk RNA sequencing (n=177 samples from n=40 patients post quality filtering) and whole exome sequencing for a subset of these samples (n=90 from n=25 patients post quality filtering). RNA and DNA from patient samples were purified from the same section to ensure high fidelity in genomic comparisons.

### RNA-Seq library preparation, sequencing, and data processing

RNA from LCM FFPE samples was isolated using the Qiagen All-Prep FFPE kit and sequencing libraries were prepared with Illumina TruSeq Paired-End Cluster Kit. Total RNA sequencing was performed on an Illumina HiSeq 2500 instrument to generate paired-end 100-nucleotide reads. After sequencing, FASTQ files for each sample were aligned to *hg38* using *STAR*^47^, and *RSEM*^48^ was used to summarize gene and transcript counts. Quality metrics were generated by *STAR* and *RSeQC*^49^ and used to evaluate sample quality. *Somalier*^50^ was used to confirm relatedness among samples from the same patient. *EdgeR 3.25.10*^51^ was used to compute normalized data by using the trimmed mean of M-values normalization and computing log2 counts per million. Gene filtering was performed using the *filterByExpr* function in *edgeR* to keep genes that had approximately 0.6 counts per million or more in at least 48 samples, with the cutoff based the median library size (17.7 million) and the histology variable (where there are 48 normal samples as the smallest group) to avoid giving preference to samples with larger library sizes. Samples were excluded (n=9 samples discarded) if they had values greater than 2 standard deviations from the mean for one or more criteria: (1) the 1st or 2nd principal components calculated using log counts per million across the filtered gene set (2) transcript integrity number median or (3) lack of relatedness among samples from the same patient as indicated by *Somalier* analysis. After sample filtering, gene filtering was recomputed as described above on the final set of high-quality samples (n=177 samples, n =21218 genes). We used *ComBat-seq*^52^ to adjust for batch effects (n=2 batches) in the Lung PCA cohort. We ranked genes to find the top 1000 genes with the highest total number of counts, then calculated the number of counts coming from those top 1000 genes for each sample. We divided this value by total counts for each sample to define an additional metric for samples that have a high percentage of their counts coming from the top 1000 genes. This metric is significantly correlated (r=-0.4, p<0.0001) to the *RSeQC* rate of uniquely mapped reads in the Lung PCA cohort, and can be computed for publicly available data sets.

### Exome DNA-seq library preparation, sequencing, and data processing

DNA from LCM FFPE samples was isolated using the Qiagen All-Prep FFPE kit and sequencing libraries constructed utilizing the Kapa Hyper Prep Kit, followed by exome enrichment with the IDT xGen Exome Research Panel v2. DNA sequencing was performed on an Illumina Novaseq 6000 instrument to generate paired-end 150-nucleotide reads. Genomic DNA from the paired normal lung tissue served as the germline control for each patient. There is no significant difference in age (t-test, p=0.69), sex (chi-square, p=0.62), tumor stage (chi-square, p=0.68), smoking status (chi-square, p=0.98), or tumor subtype (chi-square, p=0.93) between the subset of patients with both exome sequencing and RNA sequencing data compared to samples with RNA sequencing in the Lung PCA cohort. Somatic variants were identified using two different variant calling methods: *Mutect2*^53^ and *Strelka2*^54^. A mutation was called if it passed mutation calls by both algorithms. Variants in exon, splicing, and untranslated regions were assessed with a focus on exonic single nucleotide variants. Driver mutations identified by LUAD TCGA were assessed, as well as those previously identified in AAH and LUAD^10,12,55^. Each sample’s MATH score was computed as described by Mroz and Rocco^56^: 1) the mutant-allele fraction for each locus was calculated as the ratio of mutant loci to total reads, 2) the median absolute deviation was computed as the absolute difference between the mutant-allele fraction and the median mutant-allele fraction, 3) the MATH score was computed as the ratio of the median absolute deviation to the median of the mutant-allele fractions among the tumor mutated loci. Jaccard similarity was computed as the ratio of the intersection of the shared deletions, insertions, and single nucleotide polymorphisms to their union for the PML and tumor samples.

### Graph-based clustering to identify PML consensus gene modules

We sought to construct a set of gene modules designed to be robust across different LUAD PML– related data sets by adapting a method previously used to identify consensus modules that defined recurrent cancer cell states across different cancer types^57^. For each of the PML sample groups from both the Lung PCA cohort and the three additional datasets, sample and gene filtering were performed as described above for each data set independently. Each of the three additional data sets were accessed from Gene Expression Omnibus using the *GEOquery 2.51.6*^58^ package to import the gene count matrices into R: GSE102511^10^ (paired normal/PML/tumor samples from 17 patients, n= 17 AAH, n=15 normal, n=15 LUAD), GSE166720^15^ (independent samples from 50 patients with either early stage LUAD or MIA/AIS, n=17 AIS/MIA, n=33 LUAD following filtering with 2 samples removed), and GSE193725^17^ (paired normal/PML, normal/tumor, and independent samples from 42 total patients, n=23 AIS/MIA, n=10 normal, n=18 LUAD).

For each dataset, we constructed an adjacency matrix by computing Pearson correlations between gene pairs, setting negative coefficients to zero, and raising the positive coefficients to the power of six. For each individual adjacency matrix, coefficients equal or greater than the 99th percentile were set to one and the rest were set to zero. By using coefficient cutoffs based on percentiles computed for each data set, we attempted to avoid biasing the results towards data sets with higher or lower overall correlations due to technical factors. The binarized [0,1] matrices were summed, so that gene pairs with a strong correlation in multiple datasets would have values greater than 1. The summed matrix was again binarized [0,1] by setting gene–gene connections with a value of 3 or greater (the gene pair was connected in 3 or more datasets) to 1 and all other connections to 0. We then filtered out genes with few connections to other genes by removing genes with a row sum less than 10. Graph clustering performed using *infomap* clustering implemented in *igraph 1.3.5*^23^ resulted in a set of nine gene modules (n=1628 genes). Gene Ontology Biological Process 2023 gene sets and mSigDB “Hallmark” gene sets^28^ enriched (FDR < 0.05) among genes in each module were identified using *EnrichR*^25^ using *Ensembl* IDs converted to Gene Symbols with *biomaRt 2.39.4*^59^.

### Identification of archetypes

We used PMLs from four datasets to define archetypes (Lung PCA, GSE102511, GSE166720, GSE193725). The intersection of filtered genes from each dataset was used to create one gene by sample count matrix (n=137 samples, n=15968 genes). We computed gene expression residuals values for each sample using the *residuals* function in *limma 3.39.19*^60^, adjusting for study and the percentage of gene counts coming from the top 1000 genes for each sample. The residual gene expression matrix was used in all downstream analyses. GSVA^24^ was used to compute a sample score for each of the nine PML-derived modules that was used as input to the archetype analysis conducted using the *ParetoTI 0.1.13* package^26,61^ to find the positions of archetypes of a polytope (“Principal Convex Hull”) that best described the data. The archetype analysis was adapted from methods used to identify archetypes in small cell lung cancer^21^. We determined the explained variance for different numbers of archetypes, *k,* and chose the optimal number of archetypes, *k^#^*, by finding the “elbow” of the explained variance versus *k* curve. The elbow represents *k^#^*, where the explained variance starts decreasing in linear fashion when adding additional archetypes. The elbow was identified by finding the *k* most distant from the line that passes through the first (*k*=2) and the last (*kmax*=10) points in the explained variance versus *k* graph (**Supplementary** Figure 2A, red dotted line). We confirmed *k^#^* using the t-ratio metric that measures similarity between the data and the computed polytope, with a higher t-ratio indicating higher similarity. The t-ratio p-value for a given *k* (*k*=3 to *k*=6) was computed empirically as the number of times, out of 5000, that a computed t-ratio based on randomized gene modules exceeded the t-ratio. The randomized gene modules were created by randomly choosing genes (from the n=1628 gene set) to create 9 modules with identical sizes to the original modules. The lowest *k* with a significant (p<0.05) t-ratio was chosen as the optimal number of archetypes (**Supplementary** Figure 2B). Samples were scored based on their distance from each archetype (**Figure 2A**), and for each archetype, a subset of samples were assigned to an archetype. The octile (0.125) of samples closet to each archetype were assigned as primary samples of that archetype (**Figure 2B**). The 0.125 distance quantile was selected to maximize the number of samples with a primary label while keeping the fraction of samples with multiple primary labels to less than 5%.

### Characterization of archetypes

Each archetypes was annotated based on: 1) significant associations (FDR<0.05) between PML- derived gene module GSVA scores and distance from the archetype identified using linear mixed effects models where the main effects were module GSVA score and dataset and the random effect was patient, 2) significant associations (FDR<0.05) between published gene signatures^10,33^ summarized per sample using GSVA and distance from the archetype identified using linear mixed effects models where the main effects are module GSVA score and dataset and the random effect was patient, and 3) significant enrichment (FDR<0.05) of mSigDB “Hallmark” gene sets^28^ among genes with increased expression in samples closest to each archetype using GSEA with a pre-ranked gene list^30,31^. For GSEA, genes were ranked by t-statistic (using *voom*-transformed^62^ data and *limma*^60^) for their association with distance from each archetype based on linear models that included distance from the archetype, study, and the percentage of counts from the most highly expressed genes as covariates and patient as a random effect. The false discovery rate was used to adjust p-values for linear models. MIF staining and analysis was performed on samples, with cell density calculated based on the tissue area, as described and previously published by Yanagawa et al.^34^. Chi-square tests were used to determine the association between categorical archetype samples and clinical and pathologic variables (histology, dataset, smoking status, or sex). For each chi-square test contingency table, we computed an overall chi-square statistic and residuals for each cell in the table to determine which row and column combinations contributed the most to the total chi-square statistic.

### Projection of archetypes into LUAD datasets

Archetype space is continuous, so transcriptome profiles from other data sets can be placed in the polytope that defines the archetypes^21^. Additional microarray datasets (GSE68465^36^, GSE31210^37,38^, GSE30219^39^) were accessed from Gene Expression Omnibus using *GEOquery* to import the data into R, and then log2 transformed. For bulk RNA sequencing data (TRACERx, Lung PCA tumors, GSE166720), sample and gene filtering were performed as described above for each dataset. TRACERx data were downloaded from https://doi.org/10.5281/zenodo.7683605 and https://doi.org/10.5281/zenodo.760338629. For each dataset, we computed GSVA scores for the PML-derived modules on normalized gene expression data, and used these 9 GSVA scores to compute the Euclidean distance of each sample to each of the stored locations of the PML-derived archetype centers. To visualize the samples relative to the archetype positions, we used the *ParetoTI* package^26,61^ to project the stored positions of the archetypes from our PML analysis into the principal component space computed for each independent LUAD dataset.

We used several methods to determine the relationship between the archetypes and LUAD clinical and histopathologic features in each independent LUAD dataset, and in LUAD samples from the Lung PCA and GSE166720 datasets. Chi-square tests were used to determine the association between categorical archetype samples and 1) LUAD histologic patterns in the Lung PCA data and 2) tumor grade and lymph node involvement in the GSE68465 data. To determine the association between distance from each archetype center and LUAD histologic subtype (lepidic, solid, etc.), linear modeling was used with the main effect as tumor subtype and stage and the random effect as patient, where applicable (TRACERx, Lung PCA). The relationship between disease-free survival and categorical archetypes and/or distance from each archetype was examined in stage I tumors using a Cox Proportional-Hazards mixed effects model, modeling patient as a random effect in TRACERx that contained multiple samples from each patient.

## Data availability

Raw FASTQ files and associated sample and clinical data for the Lung PCA are available on the Human Tumor Atlas Network Data Portal (https://humantumoratlas.org).

## Code availability

All code will be made available upon publication.

## Acknowledgements

The authors acknowledge their dear friend, mentor, and colleague Dr. Pierre P. Massion, who dedicated his career to the discovery of biomarkers to improve the care and quality of life of his patients. Dr. Massion collaborated on the study design, patient and sample identification, and providing insights on the early phases of the analysis.

## Author Contributions

AES, SMD, MEL, SAM, and JEB obtained funding for the project. GAF, ER, MS, DPS, MM, and EJB provided pathologic review of the biospecimens. KK, GL, SZ, HL, and EG generated the data. KEA, LMT, YK, and LY processed the raw data and performed analyses. KEA created the figures. KEA and JEB wrote the manuscript. KEA, JEB, MEL, and SAM edited the manuscript.

## Corresponding Author

Correspondence should be addressed to Jennifer Beane, PhD Section of Computational Biomedicine, Department of Medicine Boston University Chobanian and Avedisian School of Medicine 72 E. Concord Street Boston, MA 02118

## Supplementary Figure Legends

**Supplementary Figure 1:**
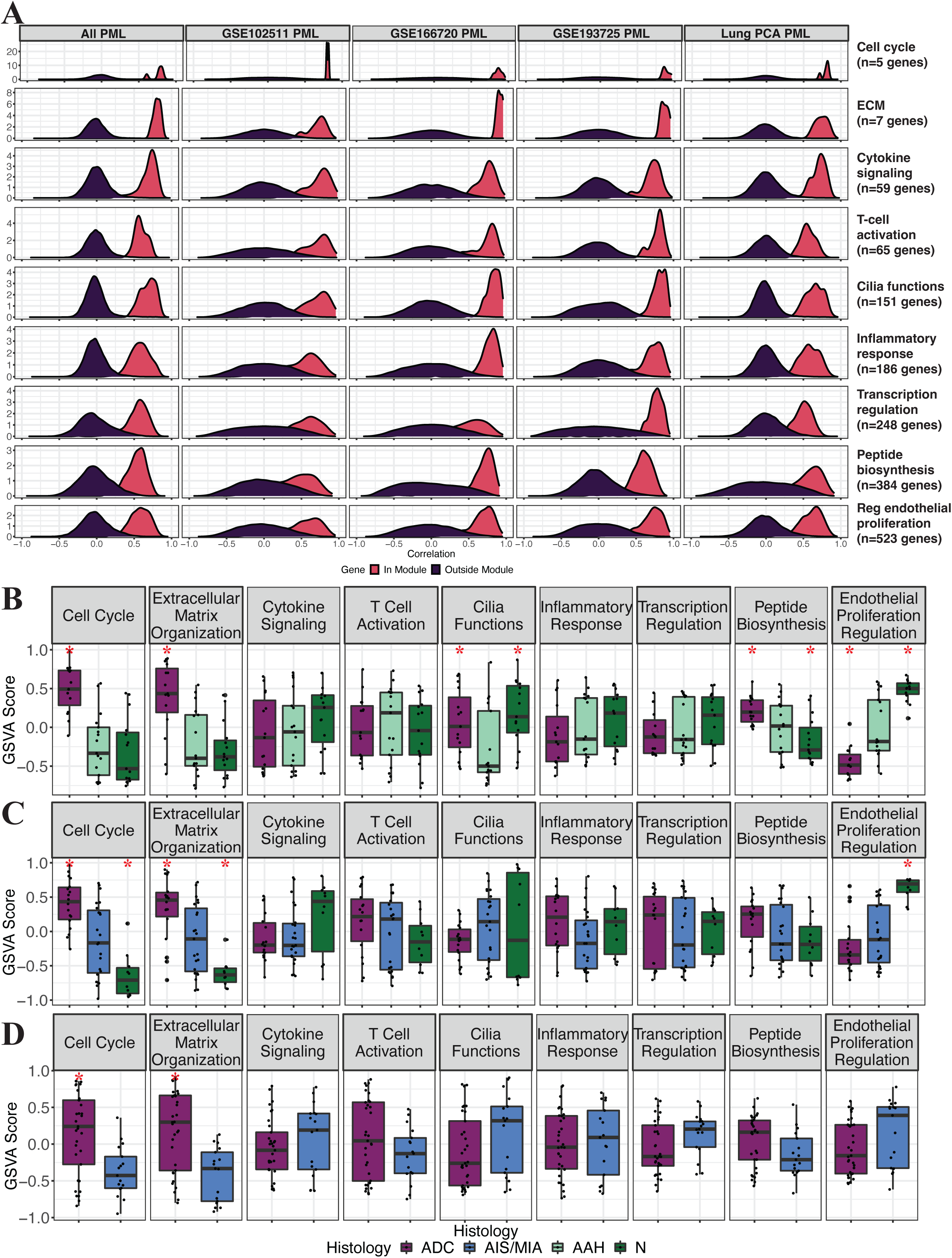
PML-derived module co-expression. (A) Density plot of correlation of each gene’s expression values to module GSVA score stratified by genes within the module and genes outside the module for all PML data (left) and PML data from each cohort (B-D) Boxplots of GSVA scores computed for each PML-derived module across (B) GSE102511 samples (n=47 samples) stratified by module and colored by histology. Asterisk (*) indicates histology is significantly different from AAH samples (linear model with patient as a random effect, p<0.05) (C) GSE193725 samples (n=51 samples) stratified by module and colored by histology. Asterisk indicates histology is significantly different from AIS/MIA samples (linear model with patient as a random effect, p<0.05) (D) GSE166720 samples (n=50 samples) stratified by module and colored by histology. Asterisk indicates histology is significantly different from AIS/MIA samples (linear model, p<0.05)

**Supplementary Figure 2:**
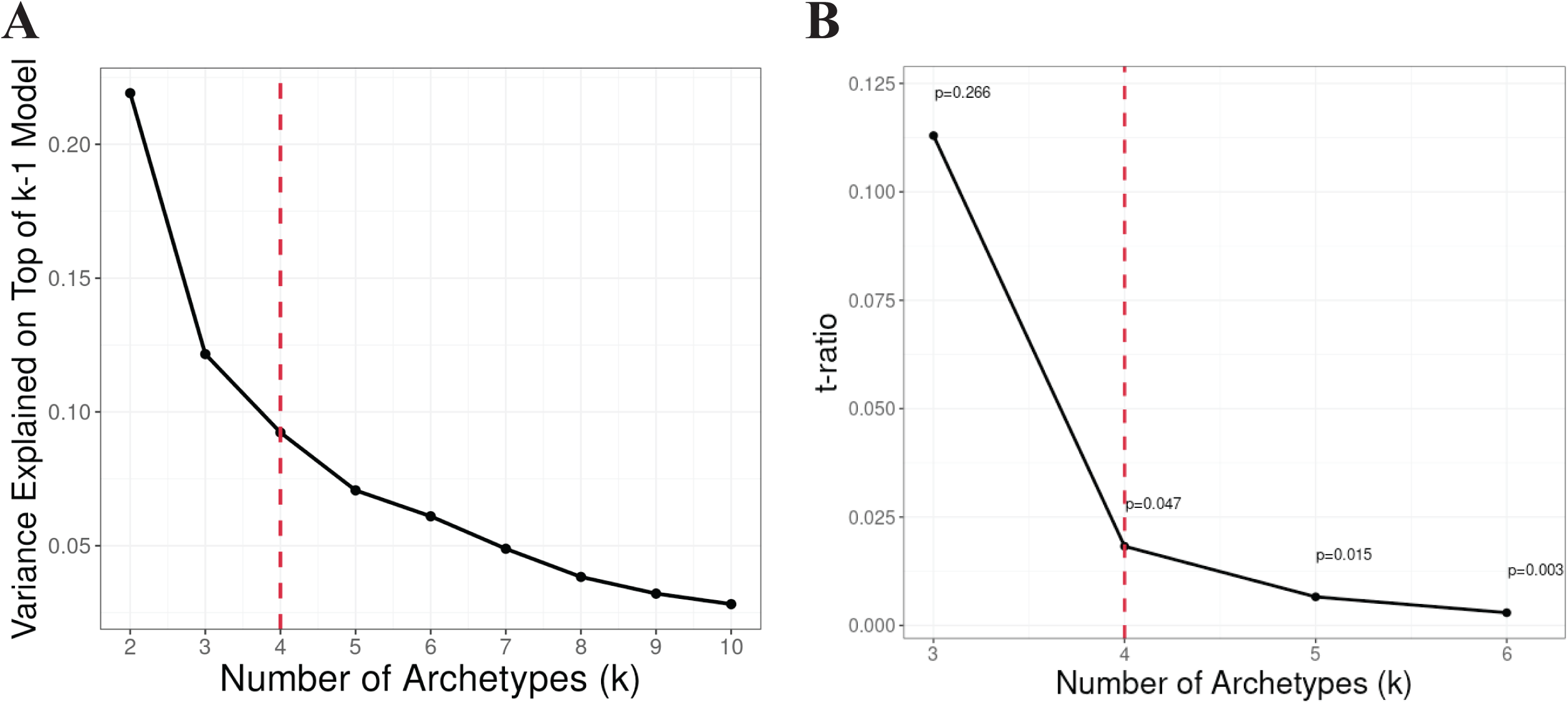
Optimization of archetype analysis. (A) Line graph of explained variance versus number of potential archetypes (*k*=2 to *k*=10), with red dashed line showing suggested number of archetypes based on elbow. Elbow computed by finding the most distant point from the line that passes through *k*=2 to *kmax*=10 (B) Line graph of t-ratio versus number of potential archetypes (*k*=3 to *k*=6) to confirm that *k*=4 is the optimal number of archetypes. We computed empirical p-values by using randomized gene modules to calculate GSVA scores. We performed this randomization 5000 times and recomputed the t-ratio for *k*=3, *k*=4, *k*=5, and *k*=6 for each iteration. Using these t-ratios as a test statistic, we determined how often the t-ratio in the randomized data was greater than the t-ratio of the PML data, and the p-value from the permutation test for each value of *k* is annotated, with a red dashed line showing the first value for *k* with a significant p-value at *k* = 4

**Supplementary Figure 3:**
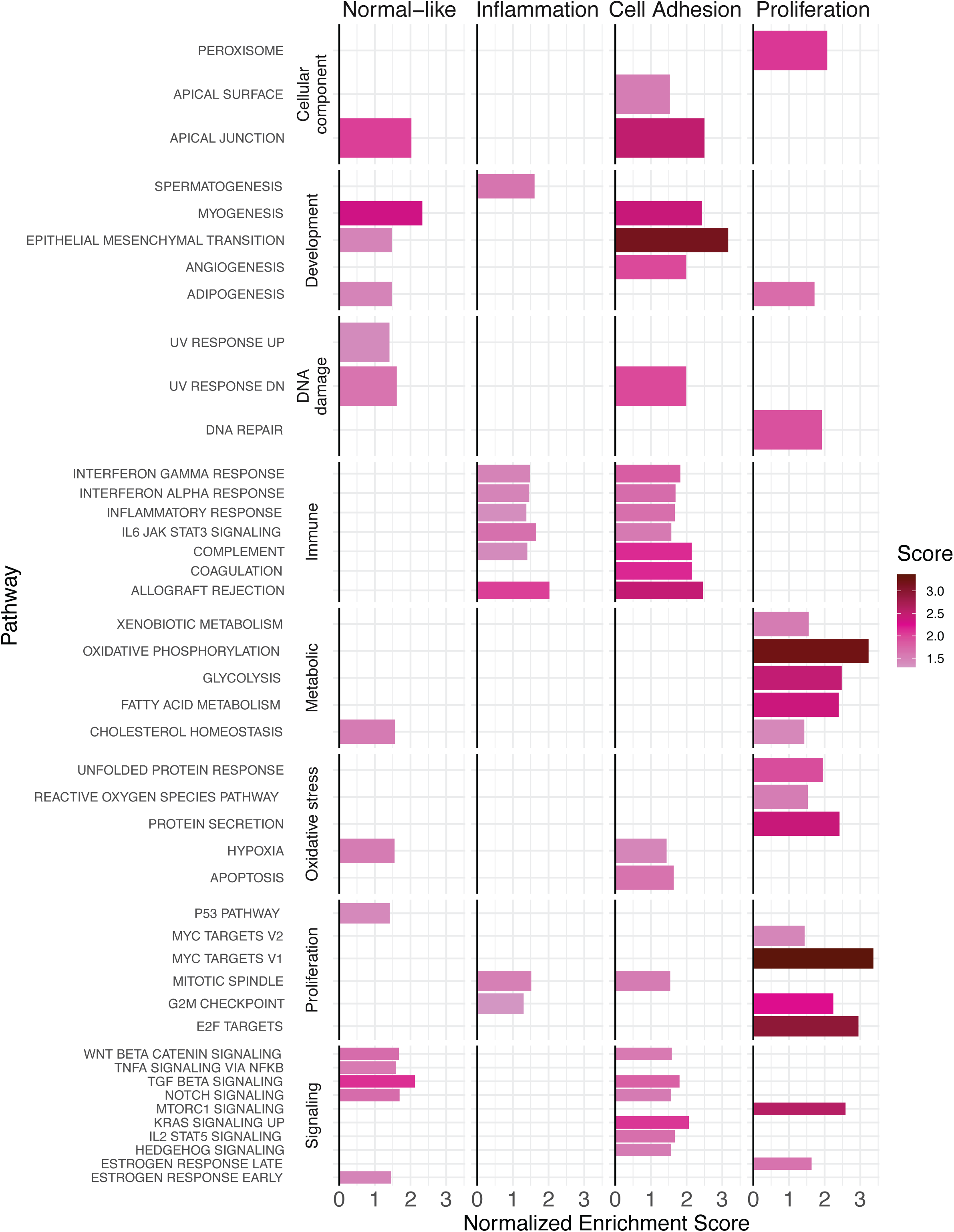
GSEA of Hallmark gene sets among genes associated with distance from archetype. Bar plot stratified by archetype and categorized by pathway type to show significant positive associations (FDR<0.05) of mSigDB ^“^Hallmark” gene sets with distance from archetypes using Gene Set Enrichment Analysis. Using limma with a model that included distance from the archetype, study, and RNA quality (Methods) as covariates and patient as a random effect, genes were ranked by t-statistic for their association with distance from each archetype. Bars are colored based on value of normalized enrichment score for each gene set, and only sets with a positive normalized enrichment score and FDR<0.05 are shown

**Supplementary Figure 4:**
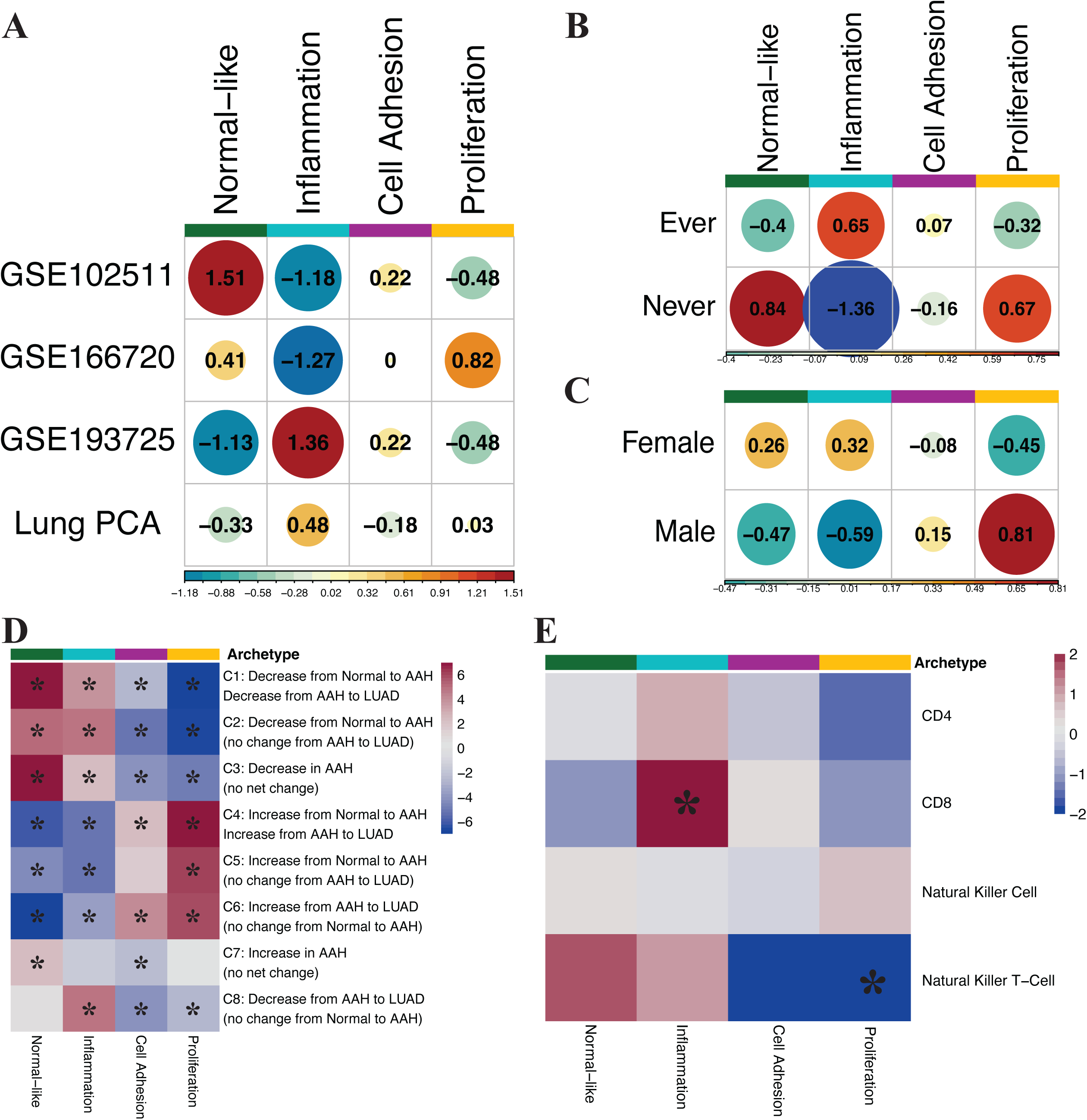
Additional characterization of PML archetypes. (A-C) Visualization of chi-square residuals from chi-square test of association of archetypes with: (A) study (n=17 GSE102511, n=17 GSE166720, n=23 GSE193725, n=80 Lung PCA PML). No archetype is significantly associated with study (p=0.33) (B) smoking status (n =17 never, n=104 ever, n=17 unknown). No archetype is significantly associated with smoking status (p=0.29) (C) sex (n =105 female, n=33 male). No archetype is significantly associated with sex (p=0.65) (D) Heatmap of t-statistics for linear model of GSVA scores for each Sivakumar et al. gene signature that differs between normal, AAH, and tumor samples modeled as a function of distance from each archetype. Asterisk (*) indicates FDR<0.05 (linear model adjusted for study with random effect for patient) (E) Heatmap of t-statistics from linear model of cell type density among select Lung PCA PML samples modeled as a function of distance from each archetype. Asterisk indicates p<0.05 (linear model with patient as random effect)

**Supplementary Figure 5:**
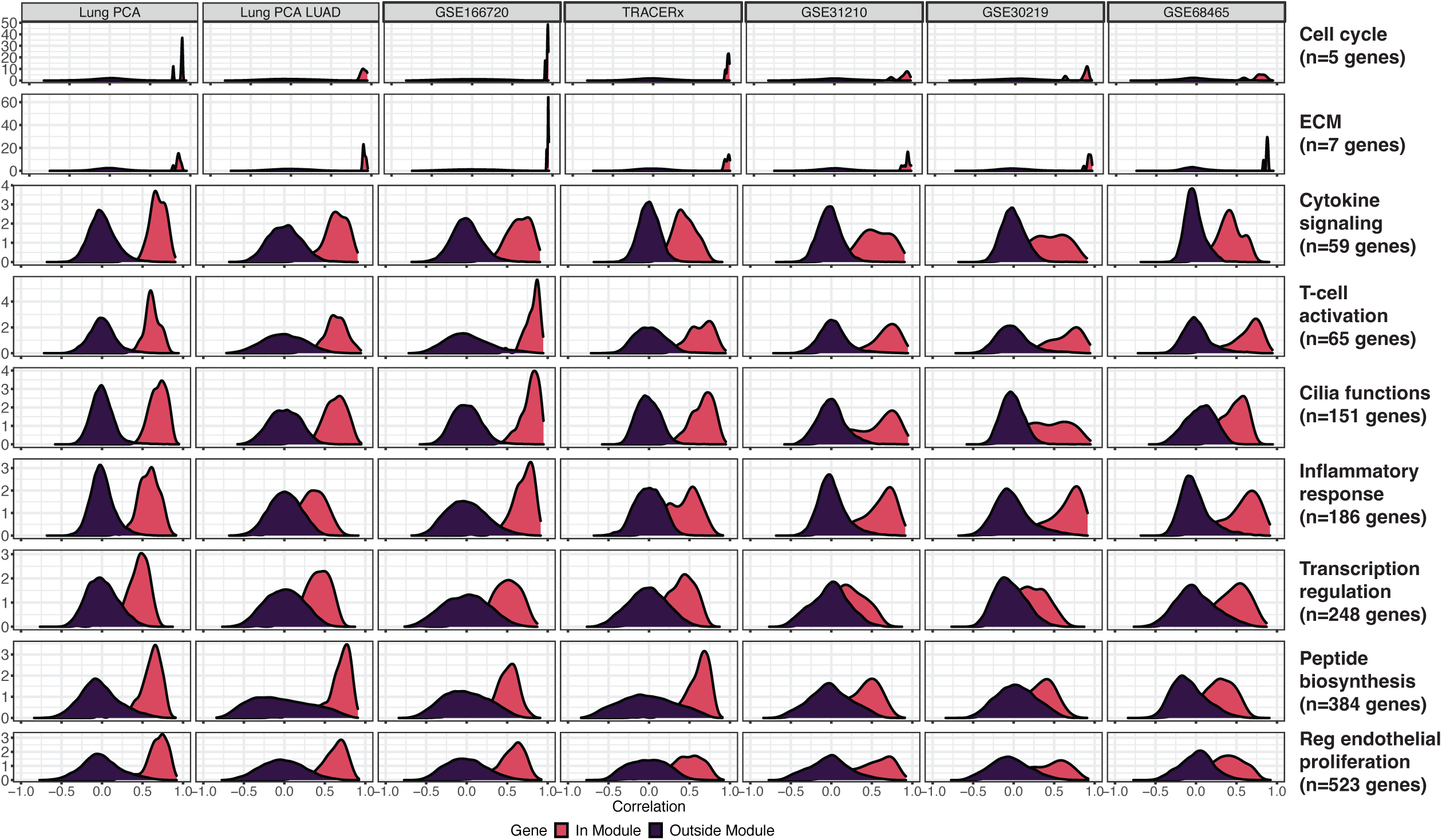
Correlation of gene expression to PML-derived module GSVA scores across LUAD datasets. Density plot of correlation of each gene’s expression to module GSVA score colored by correlation of genes within the module (pink) and genes outside the module (purple) moving across all Lung PCA samples (far left), Lung PCA tumor samples, GSE166720 tumor samples, and independent tumor data sets (moving right)

**Supplementary Figure 6:**
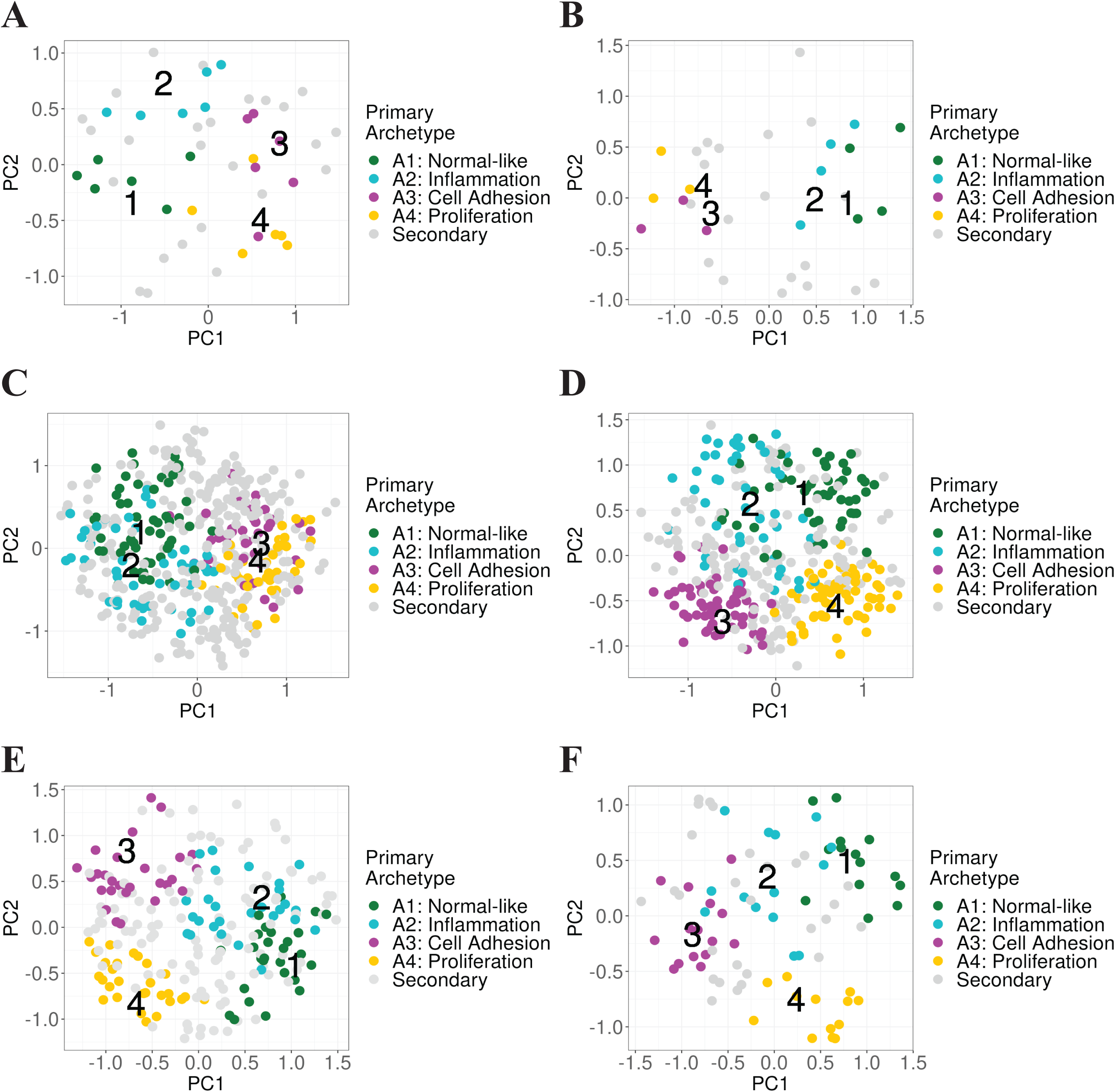
PML archetypes in LUAD cohorts. (A-F) Scatterplot of first two principal components computed from GSVA scores for PML-derived modules in LUAD datasets, with annotated projection of PML archetypes (A1-A4) into principal component space. Samples in the top 0.125 proportion of distances from each archetype were designated as primary archetypes, while other samples were labeled as “secondary”, with colors representing samples in the bin closest to archetype (A) Lung PCA tumor samples (B) Yoo et al. tumor samples (C) TRACERx tumor samples (D) GSE68465 tumor samples (E) GSE31210 tumor samples (F) GSE30219 tumor samples

**Supplementary Figure 7:**
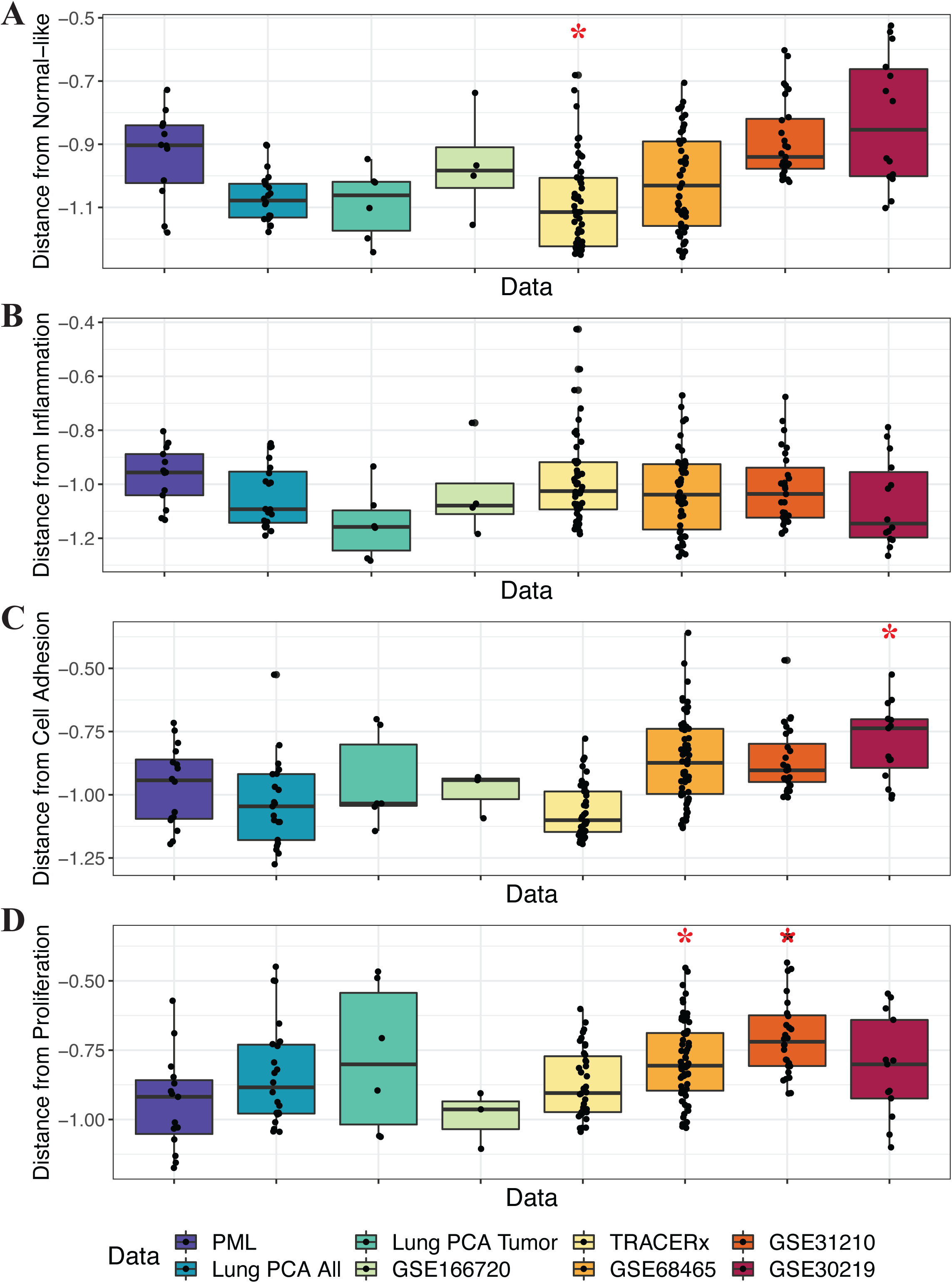
Distance between samples with categorically assigned archetypes and their respective archetype center across datasets. (A-D) Boxplots are colored by dataset, except for PMLs, which indicates all PML samples used to derive the archetypes from the 4 datasets. Samples with a categorical archetype assignment (Supplementary Figure 6) are plotted. Values closer to zero indicate the sample is closer to archetype. Asterisk (*) indicates p<0.01 (Wilcox test with distance from archetype in PML samples as basis for comparison for each data set independently). In each boxplot, distances between the samples to the archetype center are plotted on the y-axis for (A) normal-like archetype (B) inflammatory archetype (C) the cell adhesion archetype and (D) the proliferation archetype

**Supplementary Figure 8:**
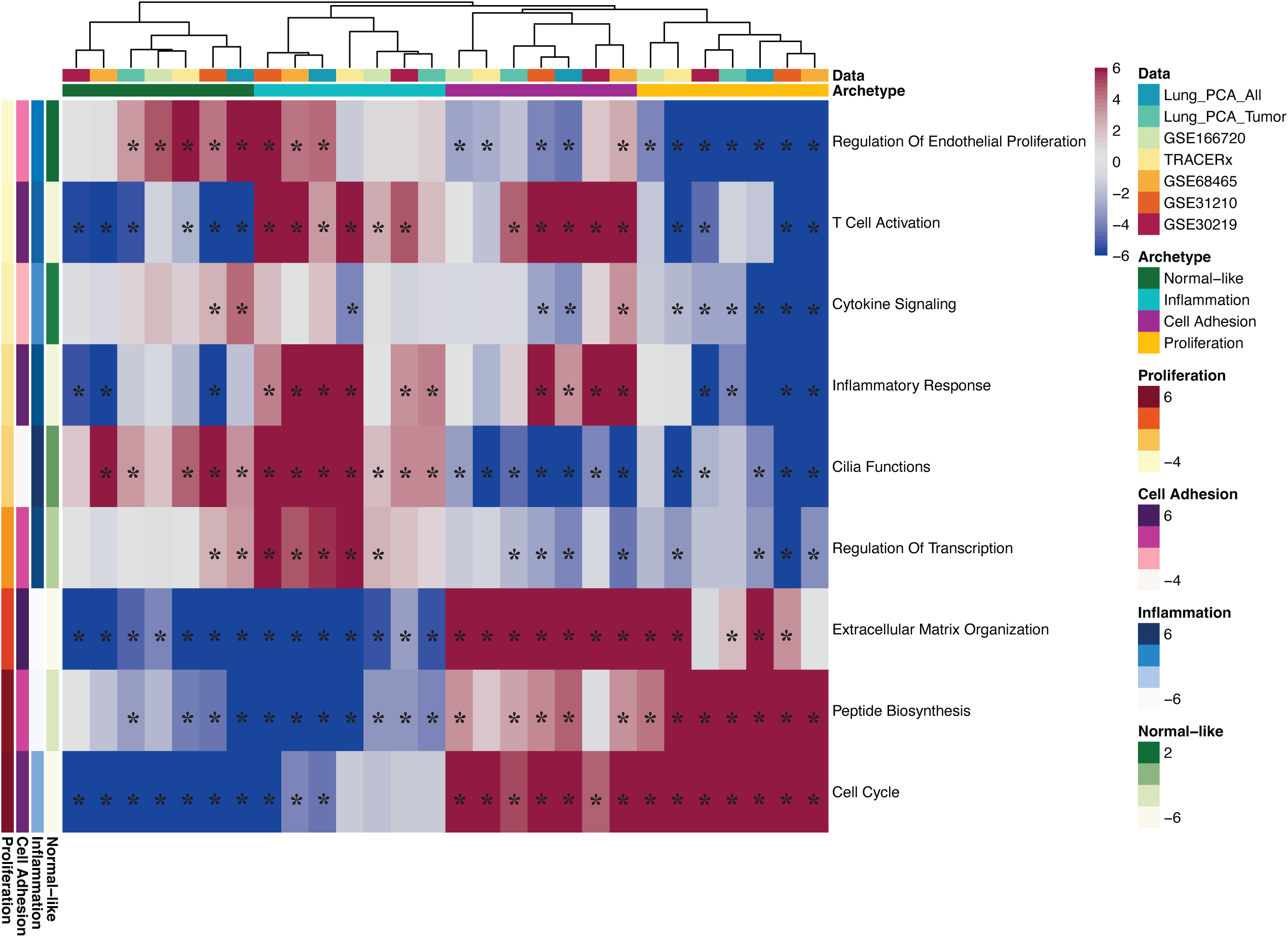
Relationship of distance from archetypes to PML-derived module GSVA scores across LUAD data. Heatmap of t-statistics from linear models of GSVA scores for each PML-derived module (rows) modeled as a function of distance from each archetype in each LUAD dataset (columns). Row annotations are t-statistics from linear models of GSVA scores for each PML-derived module modeled as a function of distance from each archetype for the PML samples (as shown in Figure 2C). Columns are semi-supervised by archetype and rows are semi-supervised by t-statistics from the proliferation archetype in the PML samples. Asterisk (*) indicates FDR < 0.05

**Supplementary Figure 9:**
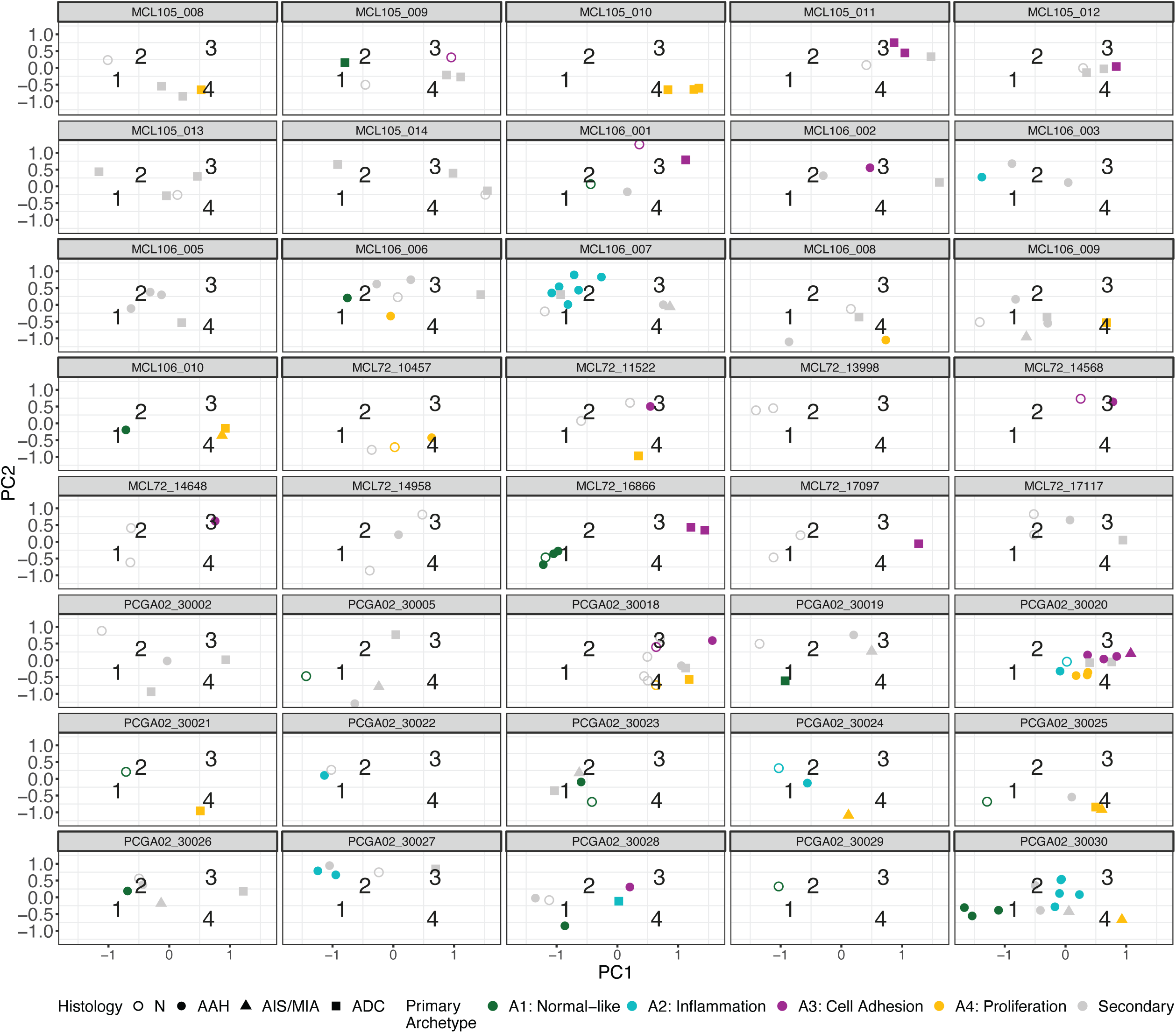
Distribution of categorical archetype samples across Lung PCA patients. Scatterplot of first two principal components computed from GSVA scores for PML- derived modules in the Lung PCA cohort. The plots are stratified by patient and samples are annotated based on their assigned archetype. Samples in the top 0.125 proportion of distances from each archetype were assigned archetypes (as shown in Figure 5A), while other samples were labeled as “secondary”, with colors representing each unique archetype. Shapes designate sample histology

**Supplementary Figure 10:**
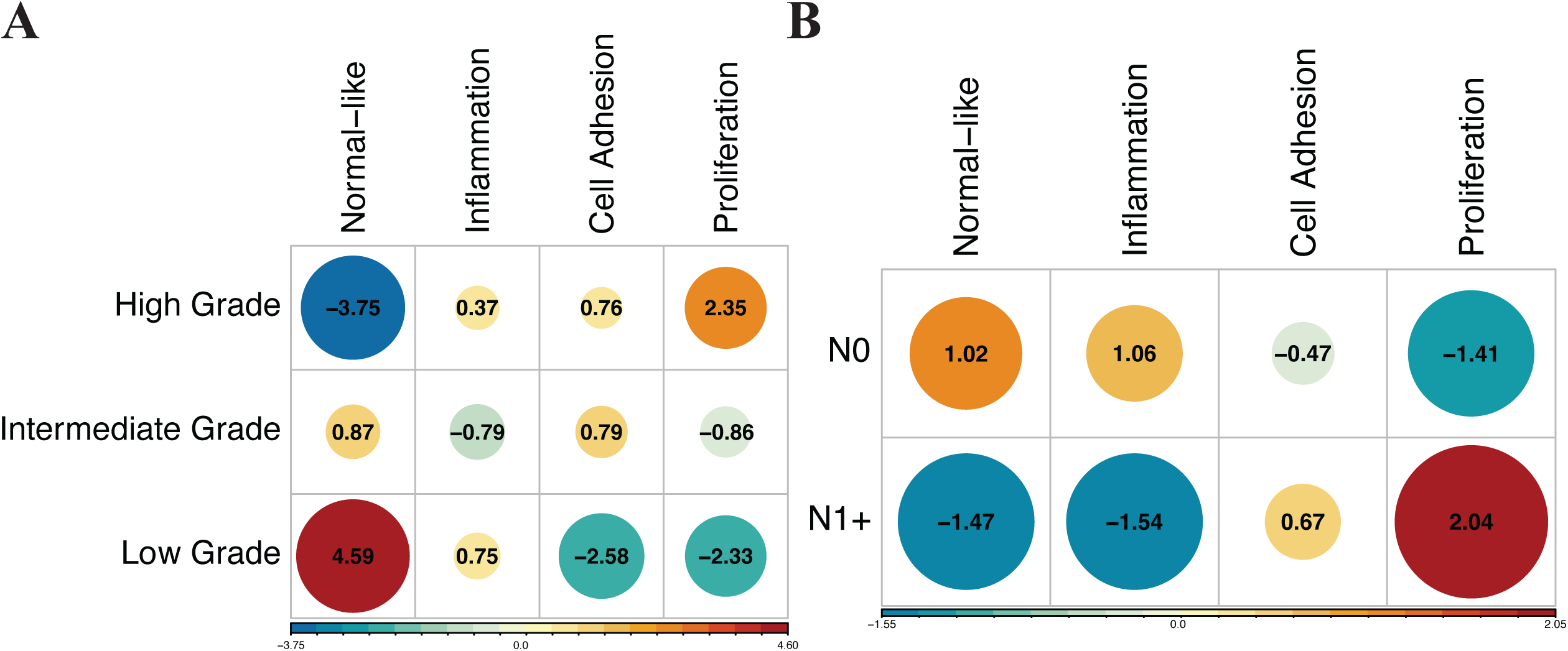
Association of archetypes with LUAD features in GSE68465. Visualization of chi-square residuals from chi-square test of association of archetypes with different disease features in GSE68465 study. Plot shows contribution of each cell to the test statistic, where circle size is proportional to cell contribution. Positive (red) values specify a positive association between the corresponding row and column variables (A) Tumor grade in GSE68465 cohort (p<0.0001) (B) Lymph node invasion in GSE68465 cohort (p=0.0038)

## Supplementary Tables

**Supplementary Table 1:** PML histology groups stratified by archetype across all PML data

|  | AAH | AIS/MIA |
| --- | --- | --- |
| Normal-like | 10 | 2 |
| Inflammation | 10 | 3 |
| Cell Adhesion | 10 | 6 |
| Proliferation | 7 | 8 |
| Secondary | 48 | 33 |

**Supplementary Table 2:** Presence of EGFR driver mutations stratified by histology in Lung PCA data

|  | Present | Absent |
| --- | --- | --- |
| AAH | 5 | 47 |
| AIS/MIA | 5 | 7 |
| ADC | 7 | 19 |

**Supplementary Table 3:**
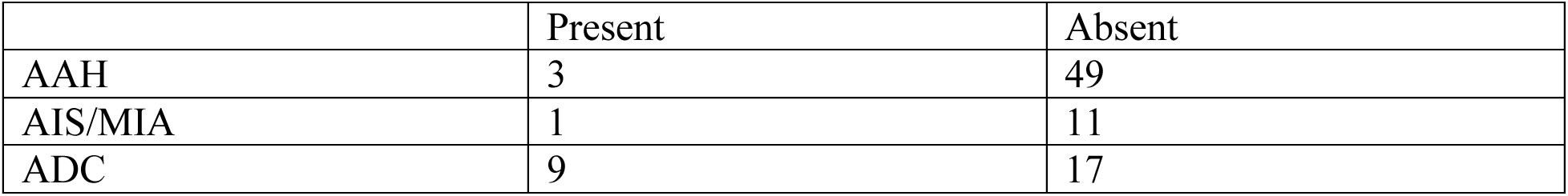
Presence of TP53 driver mutations stratified by histology in Lung PCA data

|  | Present | Absent |
| --- | --- | --- |
| AAH | 3 | 49 |
| AIS/MIA | 1 | 11 |
| ADC | 9 | 17 |

**Supplementary Table 4:** Presence of KRAS driver mutations stratified by histology in Lung PCA data

|  | Present | Absent |
| --- | --- | --- |
| AAH | 8 | 44 |
| AIS/MIA | 2 | 10 |
| ADC | 4 | 22 |

**Supplementary Table 5:** Presence of EGFR driver mutations stratified by dataset

|  | Present | Absent |
| --- | --- | --- |
| GSE102511 | 2 | 11 |
| Lung PCA | 10 | 54 |

**Supplementary Table 6:** Presence of TP53 driver mutations stratified by dataset

|  | Present | Absent |
| --- | --- | --- |
| GSE102511 | 1 | 12 |
| Lung PCA | 4 | 60 |

**Supplementary Table 7:** Presence of KRAS driver mutations stratified by dataset

|  | Present | Absent |
| --- | --- | --- |
| GSE102511 | 3 | 10 |
| Lung PCA | 10 | 54 |

**Supplementary Table 8:** Presence of JAK3 driver mutations stratified by dataset

|  | Present | Absent |
| --- | --- | --- |
| GSE102511 | 2 | 11 |
| Lung PCA | 3 | 61 |

**Supplementary Table 9:** Lung PCA histology groups stratified by archetype

|  | Normal | AAH | AIS/MIA | LUAD |
| --- | --- | --- | --- | --- |
| Normal-like | 7 | 11 | 0 | 2 |
| Inflammation | 2 | 17 | 0 | 1 |
| Cell Adhesion | 4 | 9 | 1 | 7 |
| Proliferation | 2 | 6 | 4 | 10 |
| Secondary | 33 | 25 | 7 | 29 |

**Supplementary Table 10:** Tumor grade stratified by archetype assignment in GSE68465 cohort

|  | Low Grade | Intermediate Grade | High Grade |
| --- | --- | --- | --- |
| Normal-like | 20 | 26 | 3 |
| Inflammation | 9 | 17 | 20 |
| Cell Adhesion | 1 | 29 | 26 |
| Proliferation | 2 | 22 | 35 |

**Supplementary Table 11:** Lymph node involvement stratified by archetype assignment in GSE68465 cohort

|  | N0 | N1+ |
| --- | --- | --- |
| Normal-like | 39 | 10 |
| Inflammation | 37 | 9 |
| Cell Adhesion | 35 | 21 |
| Proliferation | 31 | 28 |

